# A “ *tug-of-war*” between the NuRD and SWI/SNF chromatin remodelers regulates the coordinated activation of Epithelial-Mesenchymal Transition and inflammation in oral cancer

**DOI:** 10.1101/2024.04.05.588102

**Authors:** Roberto Stabile, Francesco A. Tucci, Mathijs P. Verhagen, Carmen Embregts, Thierry P.P. van den Bosch, Rosalie Joosten, Maria J. De Herdt, Berdine van der Steen, Alex L. Nigg, Senada Koljenović, Jose A. Hardillo, C. Peter Verrijzer, Adrian Biddle, Robert J. Baatenburg de Jong, Pieter J.M. Leenen, Riccardo Fodde

**Affiliations:** Department of Pathology Erasmus University Medical Center, Rotterdam, The Netherlands; Viroscience Erasmus University Medical Center, Rotterdam, The Netherlands; Otorhinolaryngology and Head & Neck Surgery, Erasmus University Medical Center, Rotterdam, The Netherlands; Biochemistry, Erasmus University Medical Center, Rotterdam, The Netherlands; Centre for Cell Biology and Cutaneous Research, Blizard Institute, Queen Mary University of London, UK; Immunology, Erasmus University Medical Center, Rotterdam, The Netherlands

**Author notes:** equal contributions (Francesco A. Tucci, Mathijs P. Verhagen, Carmen Embregts). current address: European Institute of Oncology IRCCS, Via Ripamonti 435, 20141 Milan, Italy (Francesco A. Tucci). current address: Department of Pathology, Antwerp University Hospital, 2650 Edegem, Belgium (Senada Koljenović).

## Abstract

Phenotypic plasticity and inflammation, two well-established hallmarks of cancer, play key roles in local invasion and distant metastasis by enabling rapid adaptation of tumor cells to dynamic micro- environmental changes. Here, we show that in oral squamous carcinoma cell carcinoma (OSCC), the competition between the NuRD and SWI/SNF chromatin remodeling complexes plays a pivotal role in regulating both epithelial-mesenchymal plasticity (EMP) and inflammation. By perturbing these complexes, we demonstrate their opposing downstream effects on inflammatory pathways and EMP regulation. In particular, downregulation of the BRG1-specific SWI/SNF complex deregulates key inflammatory genes such as TNF-α and IL6 in opposite ways when compared with loss of CDK2AP1, a key member of the NuRD complex. We show that *CDK2AP1* genetic ablation triggers a pro-inflammatory secretome encompassing several chemo- and cytokines thus promoting the recruitment of monocytes into the tumor microenvironment (TME). Furthermore, *CDK2AP1* deletion stimulates their differentiation into M2-like macrophages, as also validated on tumor microarrays from OSCC patient-derived tumor samples. Further analysis of the inverse correlation between CDK2AP1 expression and TME immune infiltration revealed specific downstream effects on CD68^+^ macrophage abundance and localization. Our study sheds light on the role of chromatin remodeling complexes in OSCC locoregional invasion and points at the potential of CDK2AP1 and other members of the NuRD and SWI/SNF chromatin remodeling complexes as prognostic markers and therapeutic targets.

## Introduction

Oral squamous cell carcinoma (OSCC) is the most prevalent type of head and neck cancer, representing over 90% of malignancies originating from the mucosal epithelium of the oral cavity (1,2). OSCC is associated with various risk factors, including tobacco and alcohol use, as well as viral infections such as human papillomavirus (HPV). However, unlike other sub-sites in the head and neck region, HPV is responsible for only a small percentage (2-5%) of OSCC cases, and its significance in this context remains uncertain (1,3). Improving the prognosis for the HPV-negative patients presents an unmet need, as mortality outcomes have shown limited improvement over the past few decades, with less than 50% of this subset of OSCC patients surviving beyond 5 years (4).

Conventional treatment modalities for OSCC, such as surgery and radiotherapy, have yielded suboptimal results and can lead to significant morbidity. Understanding the mechanisms underlying OSCC invasion and metastasis remains a major area of research as it accounts for the majority of cancer-related deaths, with more than 50% of patients experiencing cancer recurrence or developing metastases within three years of treatment (5–7).

To navigate the complex cascade of events underlying metastasis, cancer cells must acquire the ability to dynamically adapt to varying environmental conditions through reversible alterations in their cellular identity. This “so-called” phenotypic plasticity is governed by epigenetic mechanisms that do not alter the genetic code but instead control how information encoded in DNA is expressed in a tissue- and context-specific manner (6,8). In particular, chromatin remodeling complexes play critical roles in modulating gene expression programs essential for cellular homeostasis and development. These complexes facilitate dynamic changes in chromatin structure, allowing the precise regulation of gene transcription. The deregulation of chromatin remodeling is now considered a hallmark of cancer and contributes to the transient changes in gene expression patterns observed in tumor cells *en route* to their metastatic destinations (6,9,10).

The Nucleosome Remodeling and Deacetylase (NuRD) and the SWItch/Sucrose Non-Fermentable (SWI/SNF) complexes are two prominent chromatin remodelers that have been extensively studied in the context of homeostasis and cancer (11–15). They exhibit distinct yet interconnected functions and compete for binding to specific genomic loci, thereby influencing gene expression patterns. The primary function of the NuRD complex is to remove acetyl groups from histones, leading to chromatin compaction and transcriptional repression. The SWI/SNF complex instead disrupts histone-DNA contacts and mobilize nucleosomes, thereby promoting the accessibility of DNA to transcription factors and promoting gene expression (14,16,17).

In a previous study from our laboratories, the expression and functional analysis of the *CDK2AP1* gene (also known as *DOC1*, Deleted in Oral Cancer), encoding for a key NuRD subunit, has revealed its central role in the ‘tug-of-war’ between the NuRD and SWI/SNF chromatin remodeling complexes for the activation/repression of master regulators of epithelial-to-mesenchymal transition (EMT)(18). Of note, the same competition was previously shown to play a regulatory role in contexts other than cancer, namely B-cell development (19), vascular Wnt signaling (20), and the inflammatory response (21).

Here, we show that the same NuRD-SWI/SNF competition controls NF-kB activation together with other well-known inflammatory pathways and as such underlies the coordinated regulation of EMT and inflammation in OSCCs.

In the heterotypic denotation of cancer, malignant cells are embodied in their tumor microenvironment (TME) that encompasses a broad spectrum of diverse cell types, including stromal, endothelial, and immune cells, all of which play an active role in supporting and driving tumor progression (6). Immune cells such as monocytes are recruited into the surrounding extracellular matrix (ECM) through cytokines and chemokines secreted by cancer cells. Interleukin 6 (IL-6) and chemokines such as CCL2, CCL3, CCL4 and CCL5 contribute to monocyte recruitment and polarization towards tumor-associated macrophages (TAMs)(22–24). TAMs exhibit remarkable plasticity and can adopt distinct activation states, including the classically activated pro-inflammatory M1 phenotype and the alternatively activated anti-inflammatory M2 phenotype. The balance between M1 and M2 macrophages within the TME is critical for tumor development, invasion, and immune evasion (25).

Here, we show how, in OSCC, the competition between the NuRD and SWI/SNF chromatin remodeling complexes, next to the coordinated activation of EMT and inflammation, affects TME composition and in particular macrophage recruitment and polarization thus further modulating tumor behavior. The elucidation of the underlying regulatory networks and signaling pathways is likely to reveal novel therapeutic strategies centered on the NuRD-SWI/SNF axis and meant to target, in coordinated fashion, EMT in cancer cells and suppress inflammation and macrophage polarization in the TME.

## Results

### CDK2AP1 genetic ablation underlies EMT and increased invasive behavior in OSCC cancer cell lines

Previously, we reported on the key role played by the *CDK2AP1* (*DOC1*) gene in the NuRD-dependent regulation of EMT in competition with the SWI/SNF complex in the context of oral squamous cell carcinoma (OSCC)(18). Here, in order to develop new cell models to further extend on the functional analysis of the consequences of its genetic ablation, we employed the *CDK2AP1*-proficient OSCC cell lines CA1 (derived from floor of the mouth) and LM (mandibular region of the mouth) to generate several knockout (KO) clones by CRISPR/Cas9 technology (Figure 1A, left panel). RT-qPCR analysis of the EMT-related genes *SLUG* (*SNAI2*), and Fibronectin (*FN1*), previously shown to have enhanced NuRD gene promoter binding affinity (18), revealed their significant upregulation upon *CDK2AP1* ablation in both cell lines (Figure 1A, right panel). Additionally, flow cytometry analysis revealed reduced E-cadherin (ECAD) expression accompanied by the appearance of N-cadherin (NCAD) in *CDK2AP1*-KO clones when compared with the parental cell line (Suppl. Figure 1A).

**Figure 1.**
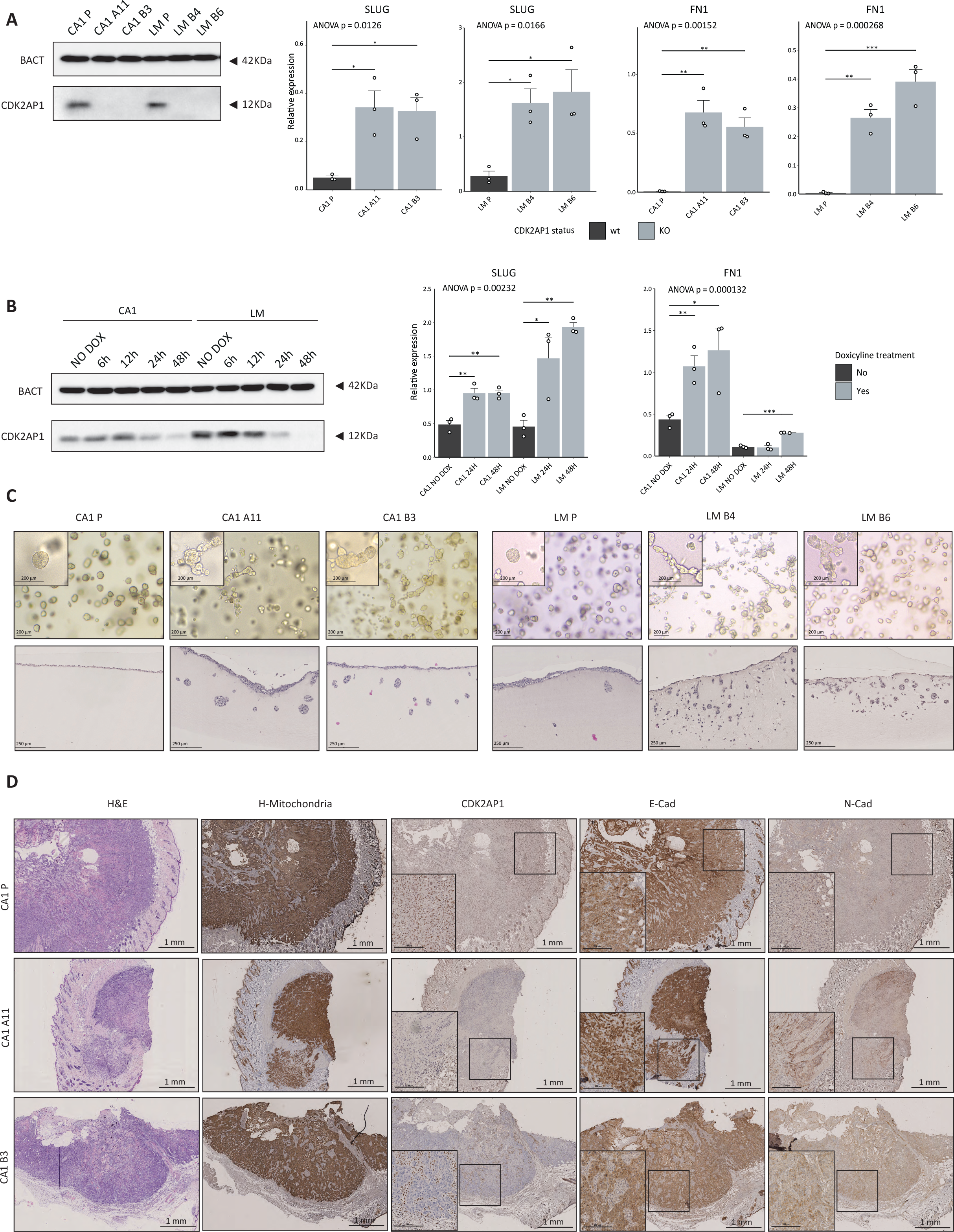
CDK2AP1 ablation in OSCC cell lines underlies epithelial to mesenchymal plasticity. **A.** Left panel: CDK2AP1 western blot analysis of our panel of OSCC parental (CA1-P and LM- P) and KO (CA1-A11 and -B3; LM-B4 and -B6) cell lines. The 12 KDa CDK2AP1 protein is observed exclusively in the parental cell lines. β-actin (BACT) was employed as loading control. The blot shown here is a representative example of 3 independent experiments. Right panel: RT-qPCR expression analysis of the EMT-transcription factor *SLUG* and the EMT marker Fibronectin (*FN1*) in CDK2AP1-proficient and -deficient cell lines. Experiments were performed in triplicate and mRNA expression was normalized to that of GAPDH. p-values denote one-way ANOVA and one-sample t-test against the Parental cell lines (*p<0.05; **p<0.01; ***p<0.001). **B.** Left panel: CDK2AP1 western blot analysis of CA1 and LM OSCC cell lines transduced with *CDK2AP1*-shRNA and induced with doxycycline (DOX) at different time points. β-actin (BACT) was employed as loading control. The blot shown here is a representative example of 3 independent experiments. Right panel: RT-qPCR analysis of the above EMT markers in *CDK2AP1*-shRNA cells at the same time points evaluated in the western depicted in **A**. Experiments were performed in triplicate and mRNA expression was normalized to that of GAPDH. p-values denote one-way ANOVA and one-sample t-test against the NO-DOX condition (*p<0.05; **p<0.01; ***p<0.001). **C.** Analysis of parental and *CDK2AP1*-KO OSCC cell lines grown in rat-tail type I collagen. The top panels depict images of cells resuspended as single cells and grown in 3D in the collagen matrix. The images were taken after 72 hours in culture to evaluate morphological differences. Higher magnifications images (20X) are shown in the top-left corner inlet. The scale bars correspond to 200 μm. The bottom panels show histological sections stained by hematoxylin and eosin (H&E) relative to the invasion assay. This was performed by seeding the cells on top of the collagen layer and let grown through the matrix. The images correspond to the 5th day of culture. The scale bars correspond to 250 μm. **D.** Immunohistochemistry (IHC) analysis of tumors obtained by subcutaneous transplantations of either parental and KO CA1 cells in NSG recipient mice. Apart from H&E (first column) the section were stained with Ab’s directed against human mitochondria, CDK2AP1, E-Cadherin (E- Cad), and N-Cadherin (N-Cad). Higher magnifications images (20X) of specific tumor areas are shown in the inlets in the bottom-left corners. Lower magnification (4X) scale bars: 1mm; higher magnification scale bars: 250 μm.

To further confirm the specific role of *CDK2AP1* in the regulation of *SLUG* and *FN1* expression, we performed knockdown experiments in the parental CA1 and LM cell lines using an inducible short hairpin RNA (shRNA)(Figure 1B, left panel, and Suppl. Figure 1B). Time-dependent RT-qPCR analysis clearly showed the progressive and direct upregulation of these target genes upon *CDK2AP1* knockdown (Figure 1B, right panel). Next, we assessed the motile and invasive capabilities of the *CDK2AP1* -KO clones. Single cells were embedded in 3D rat-tail type I collagen drops and their morphologies monitored after 72 hours. While the parental cell lines formed smooth and rounded spheroids, KO clones from both cell lines presented as collective elongated strands with numerous protrusions, indicating increased motility suggestive of pronounced invasive behavior (Figure 1C, upper panels). Subsequently, we conducted an invasion assay by seeding the cells on top of a collagen layer. Invasion events were scored after 5 days in culture (Figure 1C lower panels). H&E staining of the fixed collagen scaffolds revealed that the parental CA1 cell line predominantly grew as a monolayer on the collagen, with minimal penetration and a maximum average depth of 91 μm. In contrast, the A11 and B3 *CDK2AP1*-KO clones showed the ability to penetrate the collagen, forming islets of invading cells with average maximum depths of 283.05 and 297.02 μm, respectively (Suppl. Figure 1C). The LM clones exhibited even more extreme phenotypes, with extensive clusters of disseminating tumor cells detected inside the collagen, reaching a depth of 498.98 μm and 311.43 μm for LM-B4 and LM-B6, respectively. The invasive events of the LM parental cell line were fewer, penetrating an average depth of 137.84 μm inside the matrix (Suppl. Figure 1C).

To evaluate the consequences of *CDK2AP1* loss *in vivo*, we subcutaneously injected CA1 parental cells and the corresponding *CDK2AP1*-KO clones into immune-incompetent recipient NSG (NOD scid gamma) mice. The parental CA1-derived lesions exhibited increased growth rates when compared with their KO clones (31 days until the humane endpoint vs. 48 and 51 days in the case of A11 and B3, respectively). Increased tumor-forming efficiency was also observed with the parental CA1 line compared to the *CDK2AP1*-depleted clones (8/8 vs. 4/8 upon injections of 5x10^5^ cells). As shown in Figure 1D, immunohistochemistry (IHC) analysis of the resulting tumors showed abundant and strictly membranous expression of ECAD in the CA1 parental-derived tumors, both at the edges and in the center of the lesions. In contrast, ECAD was predominantly internalized in the cytoplasm of tumors derived from *CDK2AP1*-KO clones, particularly in the cells located along the invasive fronts. Furthermore, the *CDK2AP1*-deficient tumors exhibited areas positive for NCAD expression, indicative of a more mesenchymal-like phenotype. This observation was supported by enhanced Fibronectin (FN1) deposition by cancer cells in the *CDK2AP1* KO tumors compared to the parental-derived tumors, where the majority of FN1 staining originated from mouse cells, as indicated by the lack of human mitochondria staining (Suppl. Figure 1D).

Overall, these initial findings support EMT induction upon CDK*2AP1* ablation in OSCC.

### CDK2AP1 deletion promotes activation of pro-inflammatory pathways

To assess whether the genetic depletion of *CDK2AP1* and the consequent alteration of the NuRD- SWI/SNF competition result in downstream epigenetic modifications at genomic loci other than those involved in EMT, we conducted an extensive analysis of the ChIPseq data from the Modh-Sarip study (18).

Over Representation Analysis (ORA) of CDK2AP1-dependent NuRD chromatin binding peaks revealed an enrichment of inflammation-related pathways, and in particular of TNF-α/NF-κB (p adj < 0.01) and IL-6- JAK-STAT3 signaling (p adj < 0.01; Figure 2A), at levels comparable with those of EMT-related pathways. To validate this observation, we performed RNAseq analysis and compared the transcriptional profiles of the parental cell lines CA1 and LM with their respective *CDK2AP1*-KO clones. Principal component analysis (PCA; Suppl. Figure 2A) indicated that while the major variance component (43%), as expected, was attributed to the cell line identity (CA1 vs. LM), the impact of CDK2AP1 status was evident in the second dimension (28%). Differential expression analysis between the two groups (parental vs. KO) confirmed that *CDK2AP1* ablation triggered the dysregulation of multiple inflammation-related pathways, including interferon responses (p adj < 0.005), TNF-α signaling via NF-κB (p adj < 0.01), and IL6-JAK-STAT3 signaling (p adj < 0.005), consistent with the results of analysis of the ChIP-seq data (Figure 2B).

**Figure 2.**
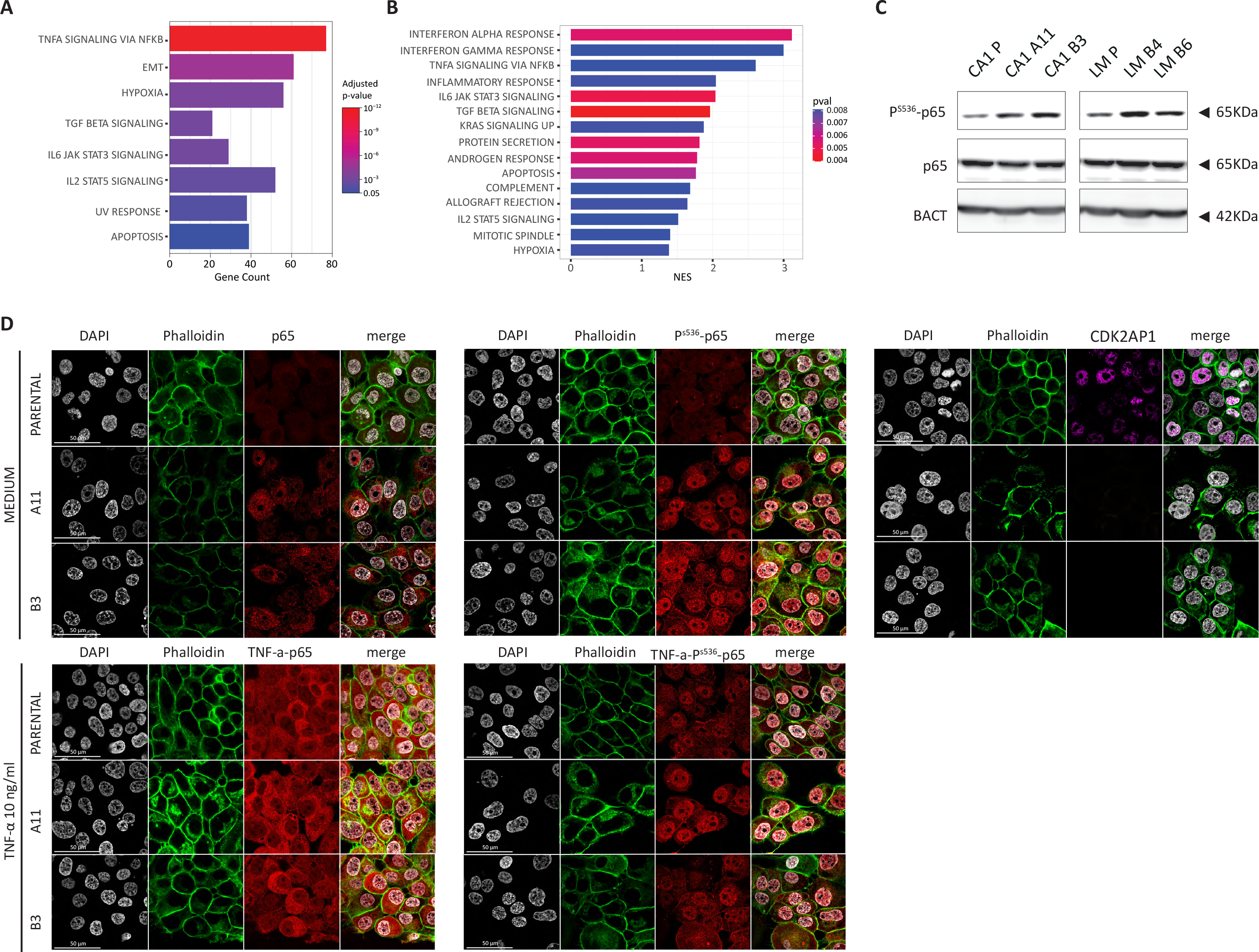
CDK2AP1 deletion, promotes inflammation via Phosphorylation of p65. **A.** Over-representation analysis (ORA) of CHIP-seq data relative to CDK2AP1-dependent NuRD peaks obtained from Modh-Sarip et al. (18). The analysis was centered on gene promoter regions (<1 Kb from the transcription start site [TSS]). Only significantly altered pathways are depicted (p adj. val <0.05).. **B.** Gene set enrichment analysis (GSEA) relative to the expression profiles obtained from parental and KO CA1 and LM cells. Significantly altered pathways (with NES >1, and pval <0.01) are shown. **C.** p65 western blot analysis in parental and *CDK2AP1*-KO CA1 cell lines. Both pan- and P- specific antibodies were employed to detect the abundance of the S536 (P -P65) phosphorylated fraction of the total. Experiments were performed in triplicate and β-actin (BACT) was employed as loading control. **D.** Immunofluorescence analysis of parental and *CDK2AP1*-KO CA1 cells culture under normal condition (upper panels) and in the presence of TNF-α for 24 hrs. (lower panels). Cells were then fixed with 4% paraformaldehyde and stained with antibodies against CDK2AP1 (purple), p65 or P -P65 (red). Nuclei and actin filaments were visualized by DAPI staining (grey) and phalloidin (green), respectively. Scale bar: 50 µm.

To identify the transcription factors (TFs) responsible for the upregulation of these pathways, we performed motif analysis of the promoter regions of the upregulated genes upon *CDK2AP1* ablation by using HOMER (26). This analysis identified several interferon regulatory factors (IRFs) and p65 as the major TFs implicated in the activation of the downstream target genes (Suppl. Figure 2C). To validate these findings, we assessed the levels and subcellular localization of the active form of p65, earmarked by serine 536 (PS536-p65) phosphorylation. Western blot analysis confirmed increased levels of PS536- p65 in the CA1 and LM *CDK2AP1*-KO clones, while the overall p65 protein expression levels remained unchanged (Figure 2C). Immunofluorescence staining further substantiated these results, demonstrating increased nuclear abundance of PS536-p65 in the A11 and B3 KO clones even under normal culture conditions (Figure 2D). Interestingly, treatment with 10 ng/ml of TNF-α for 24 hours abolished the observed differences in both p65 and PS536-p65 species, suggesting saturation of the stimulus in the KO clones (Figure 2D).

Collectively, our findings provide compelling evidence for the direct involvement of *CDK2AP1* in the regulation of inflammation-related pathways in OSCC, likely to be mediated through the activation and nuclear translocation of p65 and stimulation of IFN-γ signaling. These results underscore the multifaceted role of CDK2AP1 in both the regulation of EMT and in shaping the inflammatory microenvironment, and highlight its relevance in OSCC progression.

### The secretome of CDK2AP1 KO cells promote recruitment of PBMCs and polarization of macrophages towards a M2-like state

In view of the enhanced inflammatory profile observed upon *CDK2AP1* loss, we examined its relative clinical relevance for OSCC progression and metastasis. To this aim, we took advantage of a retrospective cohort of primary oral squamous cell carcinoma of the tongue (n=100) collected between 2007 and 2013 at the Erasmus MC and encompassing patients that received surgery as the primary form of treatment (Figure 3A)(27). As previously shown (18), the majority of tong tumors shows a mixture of CDK2AP1 positive and negative cells, with only a minority of cases demonstrating homogenous loss. Therefore, we established two categories of CDK2AP1 immunoreactivity based on the staining intensity, and an optimal threshold of 35% cancer cells negative for CDK2AP1 was found (p= 0.024; see Materials and Methods and Suppl. Figure 3A). Using this cut-off, tumor immune-infiltration strongly correlated with patients showing more than 35% of tumor cells negative for CDK2AP1 (Chi square test= 0.004; Figure 3A, right side). Based on the latter, we investigated the consequences of CDK2AP1 dysregulation on the tumor microenvironment (TME), and in particular on the recruitment of peripheral blood mononuclear cells (PBMCs) and their consequent polarization. It is known that PBMCs recruitment plays essential roles in the tumor progression and metastasis and that cancer cells are able to do so by secreting several chemokines, e.g. CCL2, CCL3, CCL4 and CCL5 among many others (28–31). Accordingly, we established the cytokine/chemokine profile of the *CDK2AP1*-KO clones in their culture media (conditioned medium or CM) by antibody arrays (LEGENDplex^TM^; see Material and Methods). Detection of the secreted active form of the chemoattractant chemokines confirmed the transcriptomic data with a significantly more pronounced chemoattractant profile in the secretome of the CA1 and LM *CDK2AP1*- KO cell lines when compared with their *CDK2AP1*-proficient parent clones (Figure 3B).

**Figure 3.**
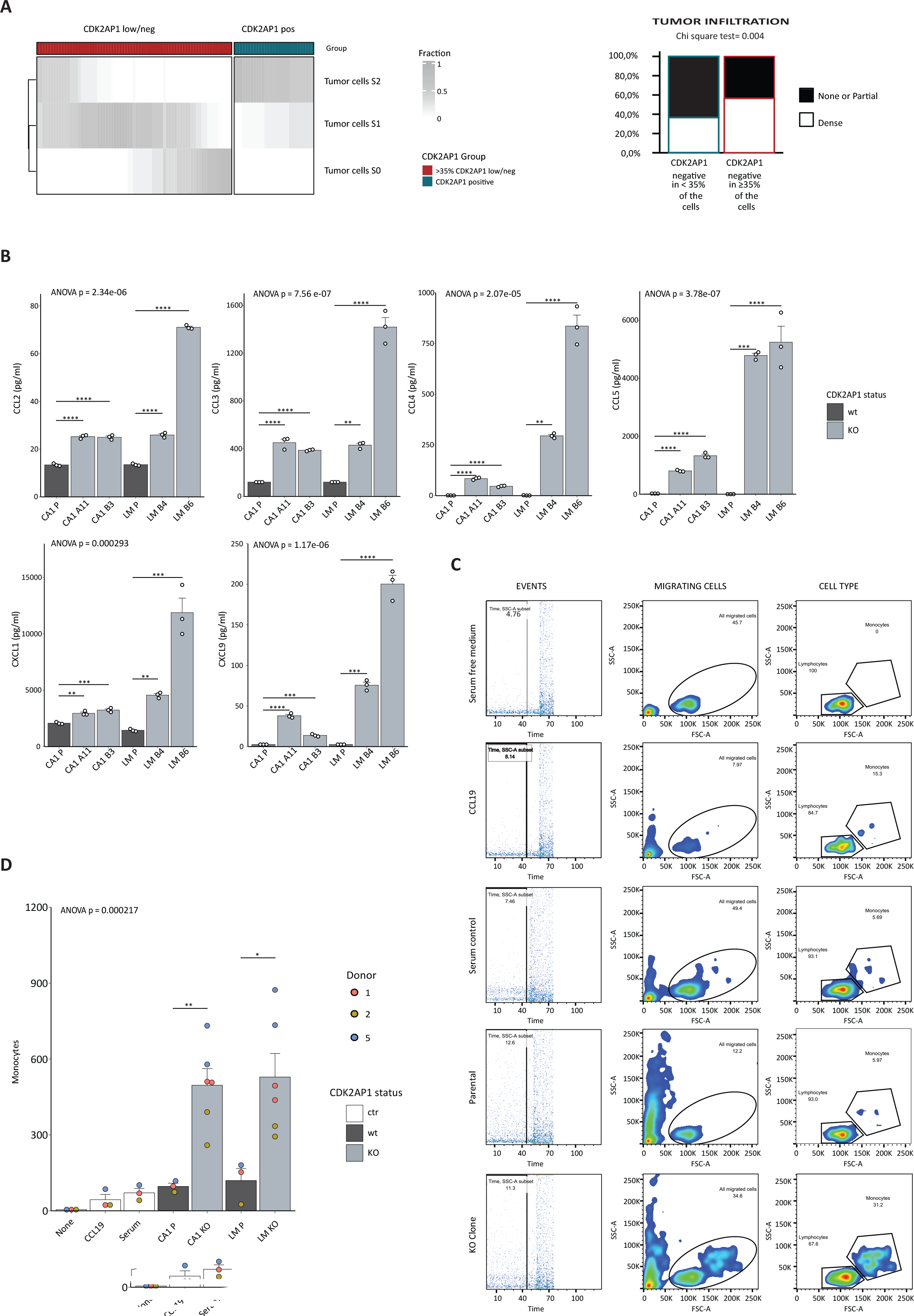
CDK2AP1 ablation results in increased immune-infiltration in vivo and PBMCs chemotaxis in vitro. **A.** Left panel: distribution of patient-derived OSCCs based on CDK2AP1 IHC staining intensities. The pre-established 35% threshold was employed to resolve low/negative from positive tumor cells (see also Supplementary Figure 3A). For each patient, the fraction of cancer cells for each staining intensity for CDK2AP1(S0, S1 and S2) is reported in grey scale. The tumors (n=100) were obtained from the RONCDOC patients cohort (for additional details see (27) and the Material and Methods section of the present study). Right panel: stacked bar plot showing the proportion of tumor infiltration (*none or partial* vs. *dense infiltration*) across the CDK2AP1 positive/low-negative patient groups. **B.** Quantification of chemotaxis-relevant cyto- and chemokines by cytometric bead assay in cell culture medium (for more details see Material and Methods). Experiments were performed in triplicate; p-values denote one-way ANOVA and Tukey test against the Parental cell line (*p<0.05; **p<0.01; ***p<0.001). **C.** *in vitro* analysis of PBMC chemotaxis. The migration of PBMCs (obtained from 3 independent healthy donors) was assessed using a trans-well assay (for more details see Material and Methods). PBMCs were allowed to migrate from the upper to the lower chamber for 3 hours. The lower chamber contained one of the following media: serum-free medium; CCL19-supplemented serum-free medium (20ng/ml); complete medium including 10% FCS; parental CA1- or LM- conditioned complete media; *CDK2AP1*-KO conditioned complete medium (CA1-A11 and -B3, LM-B4 and -B6). After 3 hrs., migrating cells were collected from the lower trans-well chamber and quantified by cyto-fluorimetry. Among the migrating events, lymphocytes were distinguished from monocytes based on their different size. Experiments were implemented in technical triplicates, with PBMCs from 3 independent healthy donors. **D.** Quantification of PBMCs chemotaxis assessed *in vitro* as described in **C**. p-values denote one-way ANOVA and Tukey test of the *CDK2AP1*-KO clones against the respective parental cell line (*p<0.05; **p<0.01; ***p<0.001).

CCL2 is one of the most prevalent cytokines expressed in the tumor microenvironment and is considered as one of the major chemoattractants of monocytes/macrophages to sites of inflammation (22,32). Upon *CDK2AP1* deletion, CCL2 secretion increased by at least 2-fold in the KO clones with maximum increase of 5.3-fold in the case of the LM-B6 clone, when compared with the parental cell lines. A similar chemo/cytokine upregulation was observed for the monocyte chemo-attractants CCL3, CCL4, CCL5, CXCL1, and CXCL9 (Figure 3B). Of note, the most extreme differences were observed with CCL5 (RANTES), a central chemokine for monocyte chemoattraction (33,34), when compared with the *CDK2AP1*-proficient cell lines: 36,5- and 60-fold increase in CA1-A11 and -B3, respectively; and a striking 1700 and 1870-fold increase in LM-B4 and -B6, respectively (Figure 3B).

Next, we challenged the ability of the different conditioned media to recruit immune cells by using a trans-well chemotaxis assay where PBMCs from 3 independent healthy donors were seeded in the upper chamber while the CM was added in the lower compartment. Serum-free media as well as unconditioned culture media, and CCL19-supplemented media were employed as controls. Migrating cells were then collected from the bottom compartments and analyzed by flow cytometry for monocytes quantification (Figure 3C). CM derived from *CDK2AP1*-KO clones attracted more efficiently monocytes from all the 3 donors. In these experimental settings, CA1 and LM KO clones recruited 5,1- (496,33 ± 65,79 vs. 96,49 ± 12,73) and 4,4-fold (528,72 ± 93,23 vs. 119,60 ± 47,74) more monocytes than their parental cell lines (Figure 3D).

Subsequently, to investigate the impact of *CDK2AP1* ablation on OSCC cancer cells in inducing macrophage polarization, M0 macrophages from CD14+ monocytes of 6 independent healthy donors were stimulated for 48 hours with various culture media, including conditioned media from our cell lines panel (see Materials and Methods). The macrophages were then examined by flow cytometry for the expression of macrophage cell surface differentiation markers: CD80, HLA-DR and PD-L1 were employed as M1-like markers, while CD200R, CD206, and CD163 were regarded as M2-like markers (Suppl. Figure 3B-C). As shown in Figure 4A, M0 macrophages stimulated to M1 with CM from parental and KO clones only showed some statistically significant differences in CD80 and PD-L1 expression, albeit at considerably lower levels when compared with those obtained upon medium complementation with the positive control stimuli IFN-γ and LPS. As far as M2-like markers are concerned, increased levels of CD200R and CD163 were observed upon stimulation with the KO conditioned media when compared with the parental CM. In the specific case of CD200R, the differences did not reach significant levels (ANOVA p= 0.1), likely due to heterogeneity among the different donors. Also in this case, the levels of upregulation obtained with the KO CM were significantly lower than those observed with the positive control stimulus IL-4. In contrast, M0 macrophages, upon stimulation with conditioned media from *CDK2AP1*-KO cells, showed a significantly increase in CD163 expression levels when compared with the CM from the parental line; this effect was similar to what observed upon stimulation with IL-10, a well-established CD163 inducer (Figure 4A; ANOVA p= 0.001).

**Figure 4.**
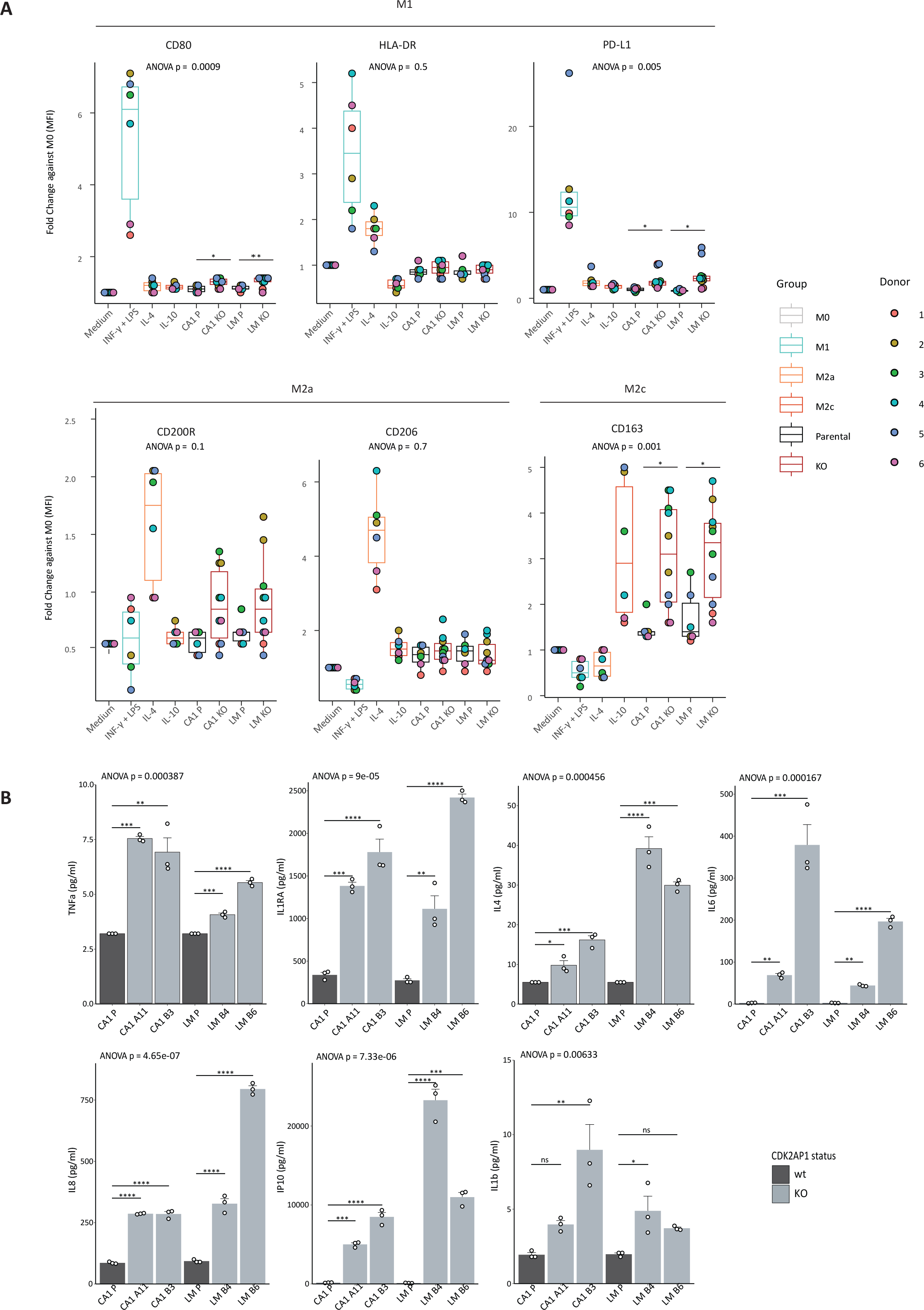
The secretome of CDK2AP1 KO cells promote polarization of macrophages towards a M2c state. **A.** Expression of M1-like (CD80, HLA-DR and PD-L1), M2-like (CD200R, CD206 and CD163) macrophage markers after 48 hours polarization with purified cytokines or with the supernatant of the parental and *CDK2AP1*-KO cell lines. PBMCs from 6 independent healthy donors were employed (see Materials and Methods). p-values denote one-way ANOVA and Tukey test of the *CDK2AP1*-KO Clones against the respective parental cell line (*p<0.05; **p<0.01; ***p<0.001). **B.** Cyto- and chemokines specific for distinct states of macroph**a**ge polarization were quantified using a cytometric bead assay in the supernatant of the culture media of the parental and CDK2AP1-KO cell lines (see Material and Methods). Experiments were performed in triplicate; p-values denote one-way ANOVA and Tukey test against the Parental cell line (*p<0.05; **p<0.01; ***p<0.001).

Analysis of the cytokine and chemokine secretions from *CDK2AP1*-KO cells, as illustrated in Figure 4B, revealed a strong statistically significant increase in inflammatory mediator production, compared with their *CDK2AP1*-proficient parental counterparts. Notably, well-established M2 polarization inducers (i.e. IL-4, IL-6, and IL-8) were elevated, with IL-6 previously recognized as a potent driver of the of M2-like polarization (22,24,35).

Altogether, these observations strongly suggest that, upon *CDK2AP1* loss, OSCC cancer cells secrete inflammatory cytokines and chemokines which efficiently recruit monocytes and polarize macrophages towards the M2-like state, with profound consequences on the tumor microenvironment in support of cancer progression and metastasis formation.

### Convergent pathways, divergent regulations: SWI/SNF loss and CDK2AP1 ablation regulate gene expression in opposite fashion

The competition between the NuRD and SWI/SNF complexes has been observed in various cellular contexts, including B-cell lineage specification and vascular Wnt signaling (19,36). Within the scope of our study, the NuRD-SWI/SNF tug-of-war plays a central role in the coordinated regulation of EMT and the inflammatory response and is as such critical for tumor progression (18,21). Hence, we set to assess whether NuRD and SWI/SNF exert opposing effects on genes involved in inflammatory pathways and the cross-talk between tumor cells and the tumor microenvironment (TME) in OSCC. To this aim, we employed inducible short hairpin RNAs (shRNAs) directed against either *BRG1* (*SMARCA4*) or *BRM* (*SMARCA2*), encoding for the mutually exclusive ATPase subunits of the SWI/SNF complex, in the CA1 parental cell line. The efficacy of the shRNAs in downregulating the protein levels of their target genes was first assessed as shown in the western Blot in Figure 5A. In view of the observed decrease in BRG1 and BRM protein expression in response to doxycycline treatment, and given their partially redundant and cooperative activity (37–40), samples were collected for RNAseq analysis after 24 hours of induction with doxycycline (Figure 5B). As depicted in Figure 5C, BRG1 knockdown had a more pronounced impact on the cellular phenotype than BRM, significantly affecting gene expression (93 genes with Log2 Fold Change > 1.5, p adj < 0.1; Suppl. Table 1), in contrast to BRM (37 genes) (Figure 5C and Suppl. Table 2). In agreement with its role in transcriptional activation, *SWI/SNF* knockdown resulted in a majority of downregulated genes (75% for BRG1 and 68% for BRM; Figure 5C). Analysis of differentially expressed genes (Log_2_ Fold change > 1.5) upon disruption of the BRM-specific SWI/SNF complex (Figure 5D, right side, and Supplementary Table 2) did not reveal any overlap with the differentially expressed genes previously identified by perturbing *CDK2AP1* expression, with the only exception of CCL5. Likewise, comparative pathway analysis of these data sets did not show any statistically significant results (data not shown).

**Figure 5.**
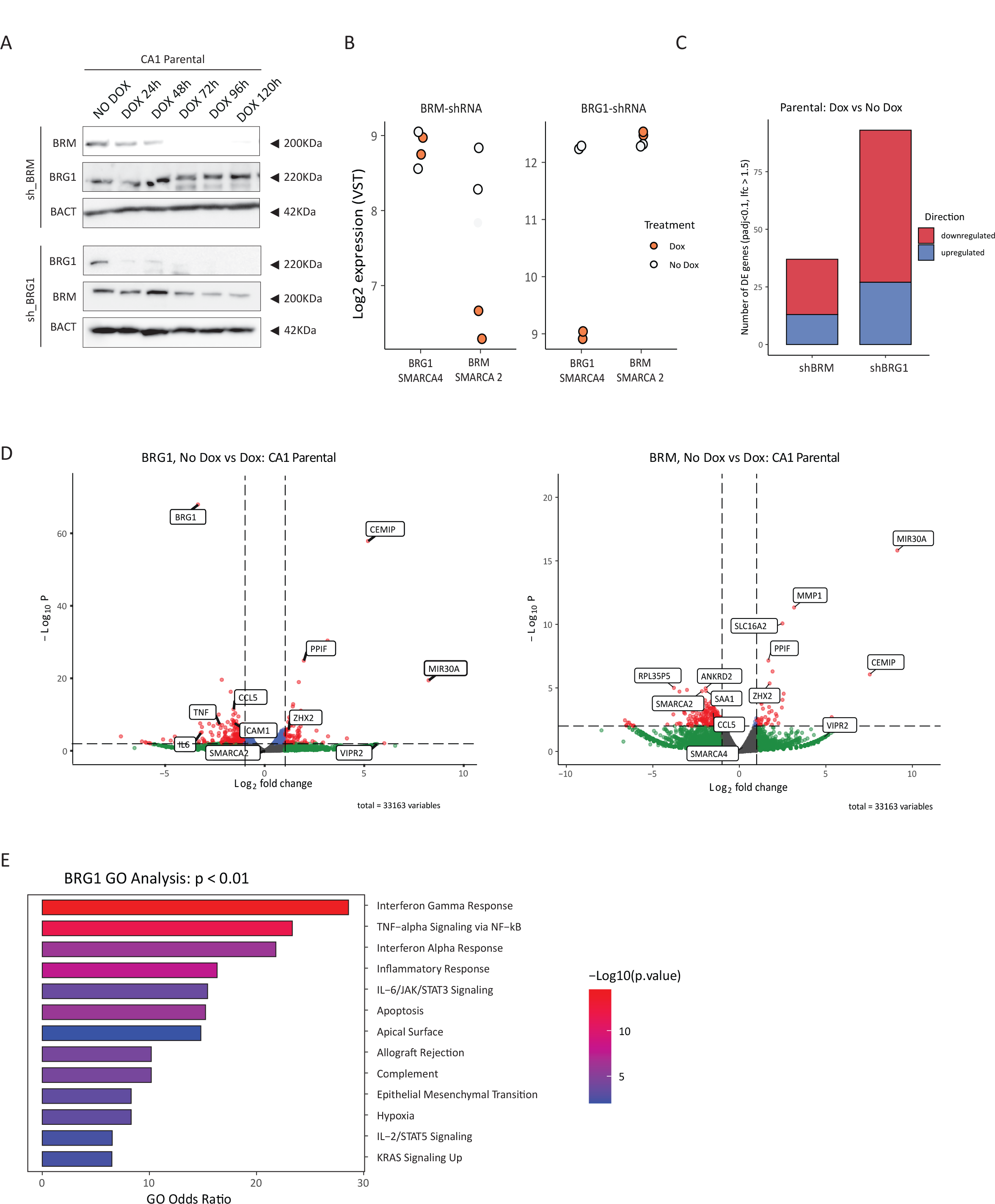
Perturbing SWI/SNF complexes, influences the same genes and pathway regulated by CDK2AP1. **A.** BRM and BRG1 western blot analysis of the CA1 OSCC cell line transduced with *BRM*- or *BRG1*-shRNA vectors and induced by doxycycline (DOX) for up to 5 days (120 hrs). β-actin (BACT) was employed as loading control. The blot shown here is a representative example of 3 independent experiments. **B.** RNAseq-based validation of the effects of the specific *BRM*- and *BRG1*-shRNAs on the expression levels of their respective *SMARCA2* (*BRM*) and *SMARCA4* (*BRG1*) targets. **C.** Overview of the differentially expressed genes upon induction of the *BRM*- and *BRG1*- shRNAs in the CA1 parental cell line. Only genes with statistically significant (p adj. <0.1 and Log Fold change>1,5) up- or down-regulated when compared with the uninduced cell line. **D.** Volcano plots relative to the differentially expressed genes upon *BRG1* and *BRM* knockdown by shRNA. Down- and up-regulated genes (log_2_ fold change> 1.5; p value <0.01) are shown on the left and right sides of each plot, respectively. **E.** Gene Ontology Pathway Analysis (GO) of the expression profiles of the CA1 Parental cell line upon *BRG1* knockdown (p value <0.01; odd ratio 2).

In contrast, differential expression analysis of the RNAseq data obtained upon downregulation of the BRG1-specific SWI/SNF complex revealed substantial overlap with the downstream effects observed upon CDK*2AP1* deletion, including genes such as *TNF-α, IL6*, and *CCL5* (Figure 3B, 4B, and 5D).

Last, gene ontology (GO) analysis of the differentially expressed genes upon BRG1-KD revealed the enrichment of the same pathways previously identified upon *CDK2AP1* genetic ablation (Figure 2A-B), including TNF-α/NFκB and IL6-JAK-STAT3 signaling, and EMT (Figure 5E).

Collectively, these results pinpoint the central role of the BRG1- and CDK2AP1-specific SWI/SNF and NuRD chromatin remodelers in the coordinated deregulation of EMT in oral cancer cells and of inflammation in their microenvironment.

### Tissue microarray multiplex immunofluorescence analysis validates the functional consequences of CDK2AP1 loss on the OSCC TME

In view of the above results, the loss of the *CDK2AP1* tumor suppressor gene in OSCC is expected to have profound consequences on the immune cell landscape of the OSCC TME. To validate our study on patient-derived tumor samples, we conducted a multiplex protein immunofluorescent staining on tumor tissue microarrays (TMAs; extensively described in our recent study (27)). Briefly, from a cohort of 141 paraffin-embedded OSCCs, three distinct areas of interest were identified in each sample, one located at the tumor center and two at its periphery. These areas were then punched from the blocks to construct TMA slides for multiplex IF analysis. Next to CDK2AP1, the tissue sections were stained with the following antibodies: CD3 (T-lymphocytes), CD14 (monocytes), CD68 (pan-macrophages), and CD163 (M2-like macrophages (25,41,42). Furthermore, we utilized the anti-cytokeratin (High Molecular Weight) antibody [34BE12], which recognizes cytokeratin 1, 5, 10, and 14, to discriminate between tumor epithelium (34BE12+ive) and the surrounding stromal tissue (34BE12-ive)(27). Computational analysis of the digital scans of the stained TMAs, implemented with the recently published SPIAT tool (43), allowed us to further improve VisioPharm analysis, not only by resolving the tumor parenchyma from the stroma, but also by establishing the composition of immune cells in the TME, their spatial distribution, and relative abundance in relation to CDK2AP1 status. Figure 6A displays 3 practical examples of the SPIAT tool capabilities to reconstruct the digital versions of the IF staining and identify and analyze the different cellular subtypes. In the first row, a core enriched by T-cells infiltration (37% of infiltrating cells) is shown. In the second and third rows, examples of high-infiltrating CD68^+^/CD163^+^ (14,3 %) and high CD68^+^/CD163^—^macrophages are depicted (Figure 6A).

**Figure 6.**
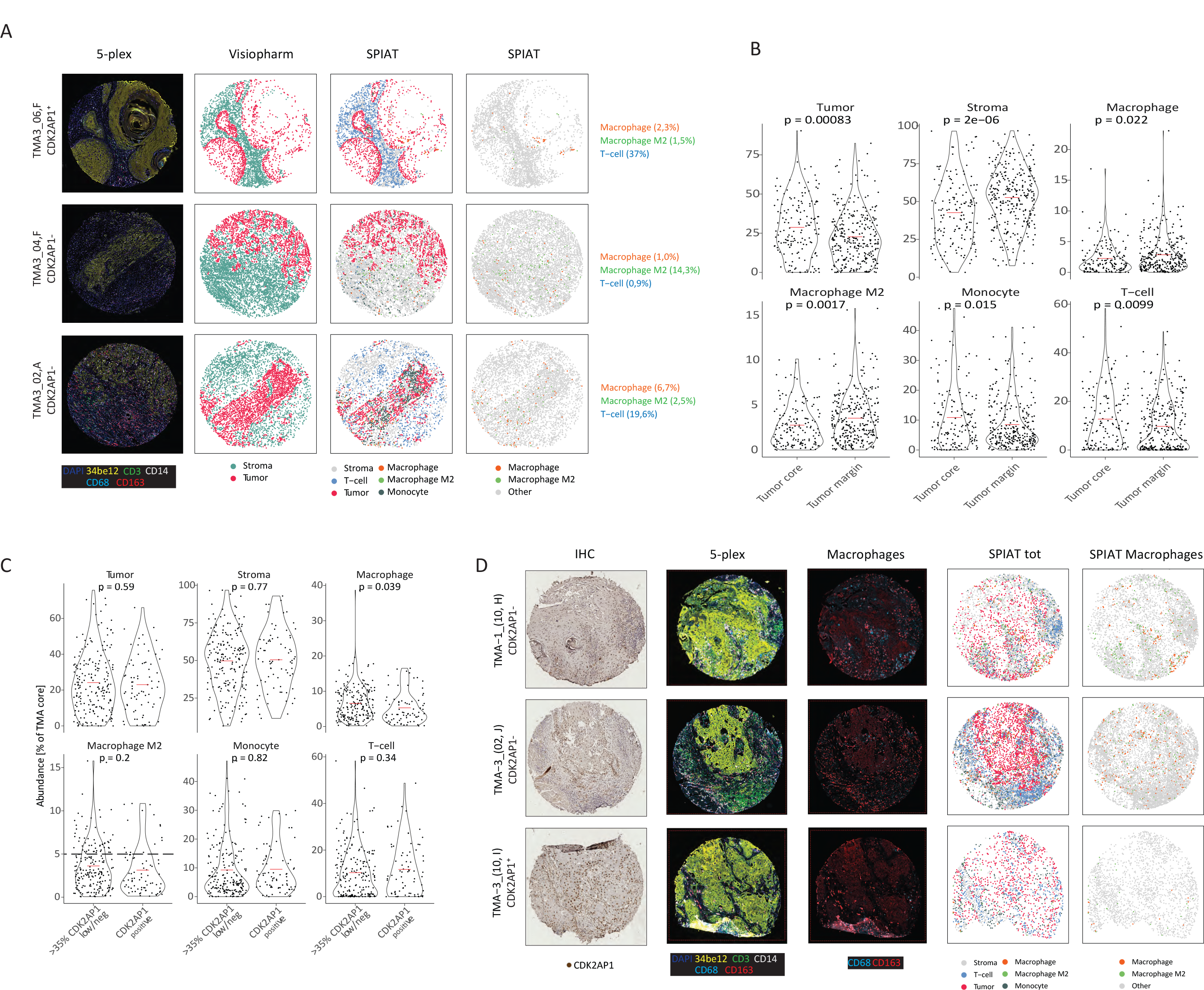
TMAs multiplex IF and ISH analysis enable improved characterization and spatial distribution of specific TMEs markers as a function of CDK2AP1 expression. **A.** Representative images of TMA fields analyzed for the presence and distribution of different immune cell markers. Next to the IF images (left column), digital reconstructions by Visiopharm and SPIAT of tumor and stromal cells are depicted. Initial Visiopharm analysis only allowed the resolution of epithelial (pink dots) and stromal (green) cells. Upon SPIAT analysis, T-Lymphocytes (blue dots), monocytes (grey), macrophages (orange), and M2c-macrophages (green) are resolved. CDK2AP1 refers to CDK2AP1 low/negative cores, whereas CDK2AP1 refers to CDK2AP1 positive cores. **B.** Violin plots relative to the differences in tumor and stromal cells, including (M2c-) macrophages, monocytes and T-lymphocytes between the tumor core and the tumor margins. P values denote paired T-Test. **C.** Violin plot relative to the same parameters analyzed in **B** across CDK2AP1-high and - low/neg cores. The dotted line in the “Macrophage M2-like” graph refers to the 5% of abundance level per TMA core. p values denote paired T-Test. **D.** Representative IHC images of CDK2AP1 protein expression in -low/neg (top) and -positive (bottom) TMA tumor cores (left column). Differences in macrophage abundance are shown by IF analysis and SPIAT digital reconstructions. CDK2AP1 refers to CDK2AP1 low/negative cores, whereas CDK2AP1 refers to CDK2AP1 positive cores. by SPIAT are depicted.

As reported in Figure 6B, regardless of CDK2AP1 status, CD68^+^ macrophages were mainly distributed in the periphery (p=0.02), while CD3^+^ lymphocytes were primarily found infiltrating the tumor core (p=0.009), as were monocytes (CD14^+^; p=0.015)(Figure 6B). When taking the CDK2AP1 staining intensity status into consideration, the relative proportion of stromal and tumor cells did not significantly change between the CDK2AP1-low/negative and positive cores (Figure 6C). However, CDK2AP1-low/negative tumors exhibited increased CD68^+^ macrophages infiltration (p=0.039; Figure 6C). Representative images of CDK2AP1-positive and -low/negative cases are displayed in Figure 6D.

As for M2-like macrophages (here defined as CD68^+^/CD163^+^), aboundance above 5% of the TMA core was only observed in CDK2AP1 low/negative tumors (Figure 6C, dotted line). However, notwithstanding this trend, these differences did not reach statistical significance when compared with CDK2AP1- proficient tumors.

In conclusion, our advanced TMA analysis confirmed the direct correlation between CDK2AP1 protein expression and CD68^+^ macrophage infiltration. The same was not found for M2-like macrophage polarization identified by CD163 expression, possibly due to technical limitations and/or other potential artifacts.

## Discussion

Although genetic alterations promote tumor initiation, cancer progression is primarily driven by epigenetic and transcriptional reprogramming (6,44,45). The ability of a cell to alter its identity in response to stimuli from the environment, also referred to as phenotypic plasticity, is advantageous over genetic alterations since it enables tumors to rapidly and reversibly respond to the plethora of volatile microenvironmental changes that cancer cells encounter along the multistep metastatic process. Accordingly, phenotypic plasticity is now widely recognized among the most clinically relevant hallmarks of cancer (6).

Here, we focused on the role of CDK2AP1 in modulating the competition between the NuRD and SWI/SNF chromatin remodeling complexes in the coordinated regulation of key downstream signaling pathways that underlie phenotypic plasticity during oral cancer progression and metastasis. Previously, we showed that post-transcriptional mechanisms including micro-RNA differential expression, rather than somatic gene mutations, underlie *CDK2AP1* loss in OSCC (27). In addition, analysis of CDK2AP1 expression in the retrospective cohort of patient-derived OSCCs revealed a strong correlation between CDK2P1 loss and poor prognosis (27). Moreover, we showed that deregulation of the competition between NuRD and SWI/SNF leads to perturbation of EMT/MET dynamics in OSCC (18). Here, extended analysis revealed that *CDK2AP1*, a gene encoding for a subunit of the NuRD complex, represents a critical player in OSCC invasion and metastasis not only because of its role in regulating epithelial-mesenchymal plasticity but also for its capacity to modulate inflammation and immunosuppression in the tumor microenvironment.

The downstream effects of the genetic ablation of *CDK2AP1* on EMT were validated both *in vitro and in vivo*. It is now widely accepted that, rather than a complete transition towards a mesenchymal identity, partial or hybrid EMT (pEMT), i.e. the co-expression of epithelial and mesenchymal genes, represents a highly metastable state that confers increased invasive and metastatic capacity to cancer cells located at the invasive front (46,47). Of note, scRNA-seq analysis of head and neck squamous cell carcinomas (HNSCC) established pEMT as an independent predictor of locoregional invasion and distant metastasis (7). From this perspective, subtle perturbations in the SWI/SNF-NuRD competition, for example through micro-RNA driven downregulation of the *CDK2AP1* gene as recently shown (27), are likely to favor the partial EMT cellular state at the invasive edge of primary oral carcinomas where they directly interact with the TME and, as shown here, trigger a pro-inflammatory state.

The coordinated activation of two hallmarks of cancer such as EMT and inflammation, the latter activated through TNF-α/NF-κB, IL-6-JAK2-STAT3 and interferon signaling, underlies phenotypic plasticity and is likely to provide a unique selective advantage to the oral squamous cancer cell during local dissemination and the formation of distant metastases.

Our studies highlight that specific alterations of the NuRD complex, and in particular those due to CDK2AP1 loss, simultaneously activate pro-inflammatory signals such as TNF-α while modulating signaling pathways with both pro- and anti-inflammatory consequences (IL-6, IL-2) to attenuate the immune responses. This apparently contradictory scenario is reflected by the chronically inflamed TME landscape where cancer cells exhibit dynamic flexibility to exploit both inflammatory and immunosuppressive signals, shaping the most appropriate niche to guarantee tumor growth, immune defense evasion and the spread of metastases (29,48).

Analysis of the cytokine and chemokine secretome activated upon *CDK2AP1* loss reveals complex scenarios where autocrine activation of NF-κB and of other inflammation-related pathways is accompanied by paracrine modification of the TME. Of note, Ramirez-Carrozzi et al. (21) previously reported on the competition between the NuRD and SWI/SNF complexes upon inflammation by showing that their antagonism controls macrophage activation upon LPS stimulation through the activation of primary (i.e. CCL5 and CCL2) and the secondary (i.e. IL-6) inflammatory response genes (21). By comparing our results with those of the Ramirez-Carrozzi study (21), a large overlap in the spectra of downstream target genes including *CCL5*, *CCL2*, and *IL-6* can be found thus pointing at the broad relevance of these cellular mechanisms in homeostasis and disease (*data not shown*).

The impact of loss or downregulation of CDK2AP1 on the TME and in particular on the recruitment of peripheral blood mononuclear cells (PBMCs) and the differentiation and polarization of infiltrating monocytes towards macrophages, was here demonstrated both *in vitro* and *in vivo*. The secretome of *CDK2AP1*-KO tumor cells exhibits an extensive chemoattractant profile for monocytes and underlies the polarization of macrophages toward an M2-like state. By taking advantage of a multiplex immunofluorescence assay tissue microarrays from a unique cohort of patient-derived OSCCs, we could validate the correlation between CDK2AP1 downregulation and the increased presence of CD68^+^ macrophages However, when it comes to tumor-associated macrophages with an M2-like profile, the overall differences with *CDK2AP1*-proficient tumors was not statistically significant, although high infiltration was only observed in *CDK2AP1*-low/negative tumors. This can most likely be attributed to technical limitations when trying to distinguish M2-like macrophages from potential artifacts by solely relying on the combination of CD68 and CD163 markers. While a growing body of evidence supports the adverse prognostic role of CD68^+^ macrophages in many cancer types (49–51), the identification of the different macrophages sub-populations in human tissues remains controversial (51–53) due to intrinsic limitations of surface marker-based analyses. While the latter are commonly used to infer functional phenotypes, the intricate nature of macrophage diversity and plasticity, influenced by complex epigenetic mechanisms as shown here, complicates this assumption (54).

Recently, two interesting reviews pointed at how novel single-cell technologies, integrated with machine learning and artificial intelligence-based analyses, will help solving this issue (55,56). Historically, macrophages were broadly classified as inflammatory (M1-like) or anti-inflammatory (M2-like) mainly based on their distinct functions and on specific markers identified through in vivo and in vitro assays. However, this classification has been challenged by the discovery of mixed expression patterns of both M1-like and M2-like genes in human tumor-associated macrophages (TAMs) across various cancer types (54,56). The advent of single-cell RNA sequencing (scRNAseq) revolutionized the field by enabling the molecular characterization of TAMs in more detail (49,51,57). These studies identified novel subsets of macrophages within tumors, taking advantage of the highest expressing genes within scRNAseq clusters for their classification. However, there remains a lack of consensus in naming the highest expressing transcripts among different studies, thus resulting in some confusion in the definition of macrophage subtypes solely based on highly upregulated gene subsets. Recent advancements have shown that macrophages comprise multiple molecular subtypes and harbor diverse functions influenced by spatial localization within the tumor microenvironment (58,59). This spatial distribution is crucial in determining macrophage functions, as different regions within the TME can induce specific signaling events that shape macrophage activation and polarization. For instance, the hypoxic core of tumors can trigger the development of angiogenic TAMs through VEGF and ANG2 secretion (60). Currently, the field is experiencing the transition from traditional M1/M2 classification to the molecular characterization of TAMs through advanced techniques like scRNA-seq and spatial transcriptomics. These advancements have paved the way for the more precise identification and targeting of diverse functional subsets of macrophages within the TME, potentially leading to improved therapeutic strategies (56).

In the near future, implementation of single-cell multiome platforms will likely shed more light on these aspects of TME biology. The recent spatial transcriptomic analysis of a small cohort of OSCC samples (61) revealed distinct profiles between tumor core (TC) and leading edge (LE). Of note, while the expression profiles characteristic of the invasive front revealed some degree of consistency across various cancer types, the equivalent profiles from the tumor cores exhibited tissue-specific characteristics. Evaluation of the prognostic impact of TC- and LE-specific gene signatures showed that high LE scores consistently associated with poor prognois and shortened disease-free survival (DFS) across multiple solid tumor types including colorectal cancer, melanoma and cutaneous squamous cell carcinoma(61). Analysis of CDK2AP1 protein expression in these tumor compartments could offer a unique perspective on how the competition between the NuRD and SWI/SNF complexes prime both the tumor cell and its surrounding microenvironment thus favoring local and systemic dissemination.

Overall, our results show how epigenetic imbalance can affect autocrine and paracrine signaling in oral cancer cells and their direct environment and elicit increased aggressiveness through secretion of pro-metastatic, inflammatory molecules. This dual impact promotes cancer development while concurrently attracting PBMCs to the tumor and inducing polarization of infiltrating monocytes into tumor-associated macrophages via inflammatory cyto- and chemokines (35).

## Materials and methods

### Cell Lines

The OSCC cancer cell lines CA1 and LM, provided by A.B., were cultured as previously described (62,63). The HEK293T cell line, obtained from ATCC, was cultured in DMEM (Thermo Fisher Scientific, #11965092) supplemented with 10% FBS, 1% penicillin/streptomycin (Thermo Fisher Scientific, #15140122), and 1% glutamine (Gibco #25030024). All cells were maintained in a humidified atmosphere at 37°C and 5% CO_2_.

### Generation of CDK2AP1 knock-out clones

Single guide RNAs targeting the CDK2AP1 gene were designed with the CHOPCHOP web tool (http://chopchop.cbu.uib.no/). The top 3 outputs, namely sgRNA1 (5’-TTCACGCTAGAGGACTGGTT-3’), sgRNA2 (5’-AAGCAAATACGCGGAGCTGC-3’), and sgRNA3 (5’-GGTGCCCCAAAGCAAATACG-3’), were cloned in the TLCV2 lenti-vector (Addgene #87360). Lentiviruses were produced by co-transfection of HEK293FT cells (Invitrogen) with the lentiviral expression vector TLCV2 with the psPAX2 (Addgene #12260) and pMD2.G (Addgene #12259) plasmids encompassing the envelope glycoprotein VSV-G (vesicular stomatitis virus G protein). Viral supernatants were collected 48-72 hours after transfection. OSCC cell lines were than infected with the lentiviruses followed by selection using 1 µg/ml puromycin (Invivogen)for 5 days. Cas9-GFP was then induced for 5 days with 1 µg/ml doxycycline. Next, GFP positive cells were single cell seeded in 96 well plate and the resulting clones were screened for CRISPR/cas9 traces in the proximity of the sgRNA by DNA sequence analysis (primers: FW: 5’- TTTGCTGAACCCATTTCTTTCT-3’; REV: 5’- ATTTTCCCCAAAAGTCTTTCCA-3’), as well as by protein detection by western blot analysis. Out of the 3 sgRNAs, sgRNA3 proved to be the most efficient for the generation of CDK2AP1 KO clones.

Generation of inducible shRNA expressing cells

In order to knockdown CDK2AP1 expression, lentiviral-inducible shRNA vectors were purchased form Horizon Discovery Ltd (Clone Id: V3THS_410413). For lentivirus production, the shRNA constructs were packaged into second-generation virus particles using psPAX2 (Addgene #12260) and pMD2.G (Addgene #12259) into HEK293T cells. Viral supernatants were collected 48-72 hours after transfection. OSCC cell lines were than infected with the lentiviruses in the supernatants and the pools were selected using 1 µg/ml puromycin (Invivogen) for 5 days. shRNA expression was induced with 1 μg/mL doxycycline at different time points. The extent of induction rate was assessed by flow cytometry while CDK2AP1 downregulation was assessed by RT-qPCR and Western Blot.

The same approach was employed to generate inducible shRNAs against BRM (Clone Id: V3THS_372090) and BRG1 (Clone Id: V3THS_317182; Horizon discovery).

### RT-qPCR and PCR analyses

Total RNA was isolated using TRIzol reagent (Thermo Fisher Scientific #15596018) and was reverse-transcribed using the high-capacity cDNA reverse transcription kit (Life Technologies #4368814), according to the manufacturer’s instructions. RT-qPCR was performed using the Fast SYBR Green Master Mix (Thermo Fisher Scientific) on an Applied Biosystems StepOne Plus Real-Time Thermal Cycling Research with three replicates per group. Relative gene expression was determined by normalizing the expression of each target gene to GAPDH. Results were analyzed using the 2-(ΔΔCt) method. Specific RT-qPCR primers here employed are listed below:

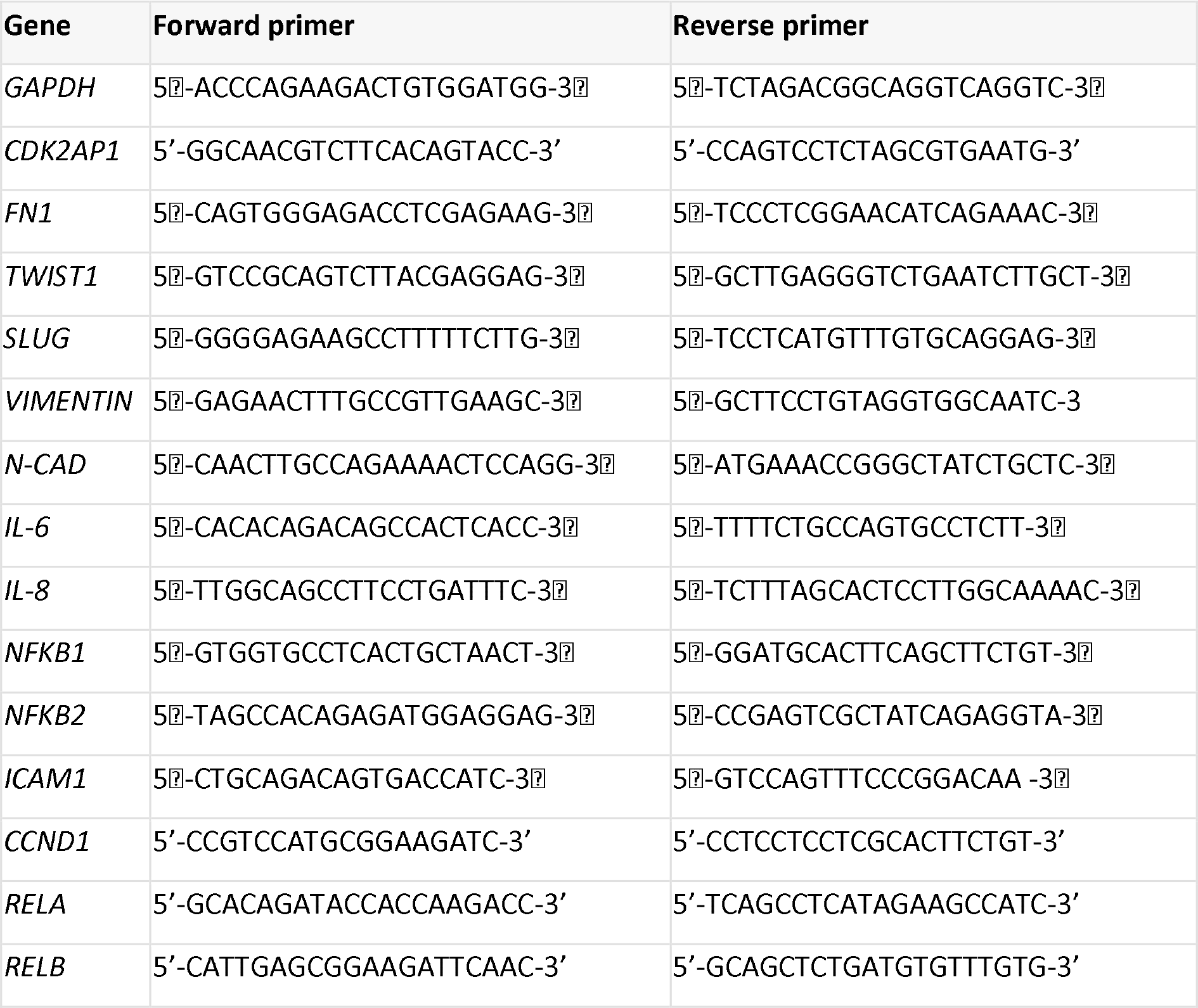

### Western blot analysis

Cells were lysed in 10 mM Tris buffer pH 7.5 containing 1% SDS, supplemented with complete™ Mini Protease Inhibitor Cocktail (Roche #11836153001), and subjected to sodium dodecyl sulfate (SDS)- polyacrylamide gel electrophoresis (PAGE). To allow detection of phosphorylated proteins, phosphatase inhibitor cocktails (Roche #04906837007) were added to the lysis buffer. Protein quantification was performed with the BCA Protein Assay Kit (Millipore); for each sample, 10 μg of protein were loaded. The membranes were incubated with primary antibodies against CDK2AP1 (1:500, Santa Cruz Biotechnology #sc-390283), β-actin (1:2000, Cell Signaling #8457), p65 (rabbit, 1:500; Abcam #ab16502), P p65 (rabbit, 1:250; Abcam #ab86299), BRM (1:1000, Cell Signaling #11966), BRG1 (1:1000, Cell Signaling #49360), followed by polyclonal goat anti-mouse/rabbit immunoglobulins horseradish peroxidase (HRP)-conjugated secondary antibodies (Dako) diluted 1:1000 for 1 hour at room temperature. The signals were detected with the Pierce ECT western blotting substrate (Thermo Fisher Scientific) using the Amersham 5 AI600 (GE Healthcare, USA).

### Immunofluorescence microscopy

Coverslips coated with a monolayer of cultured cancer cells were fixed for 151 in 4% PFA at room temperature and washed twice with PBS. Cells were first permeabilized for 15 min at room temperature with 0.1% Triton X-100 and then incubated in blocking buffer (5% milk powder in PBS-Tween) for 1 hr. at room temperature. Cells were then exposed overnight at 4°C to primary antibodies against CDK2AP1 (rabbit, 1:500; GR788 In-house production) (18), p65 (rabbit, 1:200; Abcam #ab16502), P p65 (rabbit, 1:200; Abcam #ab86299). After washing twice with PBS-Tween, coverslips were incubated for 1 hr. at room temperature in blocking buffer containing the goat anti-rabbit Alexa Fluor 568 conjugate (1:250, Life Technologies #A-11011) secondary antibodies. Cells were counterstained with DAPI to visualize the nuclei and Phalloidin Alexa Fluor 488 (1:200, #A-12379, Life Technologies) to visualize actin filaments. Coverslips were mounted in VECTAHIELD HardSet Antifade Mounting Medium, Vector Labs #H-1400) and imaged with a Zeiss LSM-700 confocal microscope. Images were processed with ImageJ (U.S. National Institutes of Health, Bethesda, MD, USA).

### Immunohistochemistry

Tissues obtained from animal experiments were fixed overnight in 4% PFA and embedded in paraffin. Paraffin blocks containing human cancer tissues were obtained from the Department of Pathology at the Erasmus Medical Center in Rotterdam. Four-µm sections were mounted on slides. IHC was performed using the EnVision Plus-HRP system (Dako) and antibodies directed against anti-human mitochondria (1:100, Merck #1273;), CDK2AP1 (rabbit, 1:500; GR788 In-house production) (18), ECAD (1:1000, Cell signaling #24E10;), NCAD (1:1000, Cell Signaling #D4R1H); FN1 (1:1000, Cell Signaling #E5H6X). Briefly, paraffin-embedded sections were dewaxed with Xylene and hydrated in 100 and 70% ethanol. Antigen retrieval was performed using pressure cooker pretreatment in Tris-EDTA buffer (pH 9.0). Subsequently, slides were incubated at room temperature in 3% hydrogen peroxidase for 10 minutes to block endogenous peroxidase activity. Tissue sections were washed and blocked with 5% milk in PBS-Tween for 1 hr. to then be incubated with the primary antibodies overnight at 4°C. Slides were washed twice with PBS-Tween and incubated with rabbit EnVision+ System HRP (K4001, Dako) or mouse EnVision+ System HRP (K4007, Dako) for 1 hour. Tissue slides were counterstained with Mayer’s Hematoxylin. Dehydration was performed by incubation in 70 and 100% ethanol followed by Xylene before slides were mounted using Pertex (Histolab #00811).

### Collagen migration assays

In order to monitor the invasion capabilities of OSCC cancer cells, parental cell lines and their CDK2AP1 KO counterparts were plated in 2 different collages models.

In the collagen droplet model, invasive behavior and morphological changes were evaluated. Briefly, 5 x 10 cells were seeded in a 20 μl droplet of rat tail collagen type 1 (4 μg/ml). Pictures were taken 4 days after plating.

In the collagen chamber model, invasive capacity was assessed using the maximum penetration measured after 5 days. Cells were seeded on top of a rat tail collagen type 1 gel (4 μg/ml) included in a scaffold at a density of 20 x 10 cells/well in 24 well plates. Following 5 days of incubation, gels were harvested, fixed in PFA 4%, embedded in soft agar, processed and sectioned for immunohistochemistry. Max distances were calculated with QuPath version 0.4.0.

### Animal experiments

All protocols involving animals were approved by the Dutch Animal Experimental Committee and conformed to the Code of Practice for Animal Experiments in Cancer Research established by the Netherlands Inspectorate for Health Protections, Commodities and Veterinary Public health (The Hague, the Netherlands, 1999). Animals were bred and maintained in the Erasmus MC animal facility (EDC) under conventional specific pathogen-free (SPF) conditions.

Sub cutaneous injections were implemented in 6- to 8-week-old NSG (NOD.Cg-Prkdcscid Il2rgtm1Wjl/SzJ) male and female mice. Briefly, 5 x 10 cells were re-suspended in 50 μl of culture medium and injected subcutaneously (4 injections in the flanks in each animal). Tumor growth was monitored by R.S. as well as the animal caretakers. Mice were sacrificed when the tumor size reached the humane endpoint. Upon collection, primary tumors were fixed, processed and scanned with the Zeiss Axioscanner 7.0 using 20x magnification.

### Chemokines and Cytokine Assays

Parental and CDK2AP1-KO cell lines were seeded at a density of 1 x 10 cells in 10 cm dishes. Supernatants were collected after 96 hours and aliquots were stored at -80°C. Chemo- and cytokines concentrations were determined using the Legendplex bead assays (BioLegend). For this project, LEGENDplex™ Human Macrophage/Microglia Panel (13-plex) and LEGENDplex™ HU Proinflam. Chemokine Panel 1 (13-plex) were used. Bead analysis was performed by a BD FACS Lyric flow cytometer and the data was analyzed using FlowJo V10.6.2.

### PBMCs chemotaxis

Peripheral blood mononuclear cells (PBMCs) were isolated from buffy coats obtained from healthy blood donors (Sanquin). Written informed consent for research use of donated blood was obtained by the Sanquin blood bank. Peripheral blood mononuclear cells were obtained by density centrifugation using Ficoll Paque PLUS (GE Healthcare). Migration of PBMCs from 3 donors was assessed using 24-well transwell chambers, where the two compartments were separated by a 5 μm pore size polycarbonate membrane (Corning, #3421). The upper compartment contained 100 μm of PBMCs cell suspension (1 x 10 cells/well) in RPMI-1640 serum-free medium. The lower compartment contained 500 μl of either cancer cell line derived conditioned medium or control media (serum free medium; 2% serum control; CCL19 (20 ng/mL, R&D Systems). The PBMCs were allowed to migrate from the upper chamber to the lower one for 3 hours. At the end of the assay, the upper well was removed and the cells in the lower compartment were analyzed with a cytofluorimeter. Migrating cells were counted distinguishing lymphocytes from monocytes based on size differences.

### Macrophage maturation and polarization

Monocytes were obtained from PBMC fractions from 6 healthy donors by magnetic associated cell sorting using CD14 beads, following manufacturers guidelines (Miltenyi Biotec). Purity of the sorting was confirmed by flow cytometry using a BD Lyric flow cytometer (BD Biosciences).

Monocytes were seeded at a density of 1 x 10 cells per well in 96-well plates and incubated to mature for 6 days in RPMI-1640 medium containing 10% pooled human serum (Sanquin), 1% (v/v) GlutaMAX (Gibco), and 20 ng/mL monocyte colony-stimulating factor (M-CSF, R&D Systems) at 37°C with 5% CO_2_. Medium was replaced on day 2 and 4.

Mature macrophages were stimulated for 48 hours with complete medium containing IFN-γ (20 ng/mL, R&D Systems) and LPS (100 ng/mL, Sigma Aldrich), with IL-4 (20 ng/mL, R&D Systems), or with IL-10 (20 ng/mL R&D Systems), to induce the M1- and M2- phenotype, respectively. To investigate the effect of CDK2AP1-KO cancer cells on polarization, macrophages were stimulated with their conditioned medium as well as with the control medium from the CDK2AP1 parental lines. Expression of macrophage phenotypical markers was investigated by flow cytometry. Non-polarized M0 macrophages, cultured for 48 h with complete medium without additional cytokines, were taken along as controls for each donor. Macrophages were dissociated from the wells using Accutase (Merck Millipore) and washed twice with PBS before staining for 30 min with the fixable viability dye ZombieViolet (Biolegend). Cells were fixed with 4% PFA for 15 min, and after Fc receptor blocking with Human TruStain FcX (Biolegend), cells were stained with the following antibodies in FACS buffer (PBS with 2% fetal calf serum, 0.2 mM EDTA, 0.01% sodium azide): anti-CD80-FITC, anti-HLA-DR-APC-Cy7, anti-PD-L1-APC, anti-CD163-PE, anti-CD200R-PE- Cy7, and anti-CD206 BV786 (all from Biolegend). Mean fluorescent intensities were quantified by flow cytometry using a BD Lyric flow cytometer (BD Biosciences) and the data were analyzed using FlowJo V10.6.2. Cell culture supernatant was collected 96 h after seeding 1 x 10 cells in 10 cm dishes and stored at −80°C. Aliquotes of the same media were used for quantification of cytokines using the Legendplex cytometric bead assay (Biolegend) as described above.

### Patient cohort, immuno-histochemistry and tumor tissue microarrays

Tissues from 100 primary oral squamous cell carcinomas of the tongue surgically removed between 2007 and 2013, were obtained from the tissue bank of the department of Pathology of the Erasmus Medical Center and recorded within the framework of the RONCDOC project, an initiative undertaken by the RWHHT (Rotterdamse Werkgroep Hoofd-Hals Tumoren). The cohort includes tongue squamous cell carcinoma (TSCC) removed by surgery as primary treatment at the Erasmus MC Cancer Institute. Cases with a previous history of head and neck cancer were excluded.

For all subjects included in this cohort, patient characteristics, comorbidity, TNM staging, treatment protocol, histopathological characteristics, recurrent disease and survival have been recorded. Tissue preparation and staining strategy were performed as described in Herdt et al.(64). Briefly, using a microtome, consecutive 4 μm sections were cut from the formalin-fixed paraffin-embedded (FFPE) cancer tissues. Hematoxylin & eosin (H&E) as well as CDK2AP1 immunohistochemical staining (1:250, Santa Cruz #sc-390283) were evaluated and scored by a pathologist. The obtained data were then correlated with the clinical-pathological information included in the cohort.

The TMAs were obtained from the same FFPE blocks. Three regions of interest for each tissue block (1 in the center of the tumor, 2 in the tumor border) were selected for sampling (1 mm diameter each) and included in the TMAs blocks.

### TME analysis by multiplex immunofluorescence

Multiplex immunofluorescence (IF) analysis was performed by automated IF using the Ventana Benchmark Discovery ULTRA (Ventana Medical Systems Inc.). Four-µm formalin-fixed paraffin-embedded (FFPE) sections were stained. In brief, following deparaffinization and heat-induced antigen retrieval with CC1 (#950-224, Ventana) for 32 min., anti-CD68 was incubated for 20 min. at 37°C followed by omnimap anti-mouse HRP (Ventana #760-4310) and detection with DCC (Ventana #760- 240) for 8 min. An antibody denaturation step was performed with CC2 (Ventana #950-123) at 100°C for 20 min. Secondly, incubation with anti-34BE12 was performed for 4 min at 37°C, followed by omnimap anti-mouse HRP (Ventana #760-4310) and detection with R6G (Ventana #760-244). An antibody denaturation step was performed with CC2 (Ventana #950-123) at 100°C for 20 min. Third, anti-CD163 was incubated for 32 minutes at 37°C, followed by Omnimap anti-mouse (Ventana #760-3210) and detection with Red610 (Ventana #760-245). An antibody denaturation step was performed with CC2 (Ventana #950-123) at 100°C for 20 min. Fourth, anti-CD14 was incubated for 32 min. at 37°C followed by omnimap anti-rabbit HRP (Ventana #760-4311) and detection with Cy5 (Ventana #760-238) for 8 min. An antibody denaturation step was performed with CC2 (Ventana #950-123) at 100°C for 20 min. Last, anti-CD3 was incubated for 32 min. at 37°C followed by omnimap anti-rabbit HRP (Ventana #760-4311) and detection with FAM (Ventana #760-234) for 8 min. Finally, slides were washed in phosphate-buffered saline and mounted with Vectashield containing 4’,6-diamidino-2-phenylindole (Vector laboratories, Peterborough, UK). Slides were imaged with Axioscan Zeiss.

| Antibody | Dilution | Species | Company | Clone | Incubation time |
| --- | --- | --- | --- | --- | --- |
| CD3 | 2,05 ug/ml | Rabbit | Ventana | 2GV6 | 32 min. |
| CD14 | 1,04 ug/ml | Rabbit | Atlas Antibodies | polyclonal | 32 min. |
| CD163 | 2,14 ug/ml | Mouse | Cell Marque | MRQ-26 | 32 min. |
| CD68 | 3,49 ug/ml | Mouse | Ventana | KP1 | 20 min. |
| 34BE12/Cytokeratin | 4,05 ug/ml | Mouse | Cell Marque | 34BE12 | 4 min. |

### TMAs bioinformatic data analysis

Three tissue micro arrays (TMA) were constructed with punches from the tumor core and periphery, from a cohort of OSCC patients (N = 462 cores, N = 164 tumors). TMAs were cut to a thickness of 4 µm and stained with antibodies directed against Cytokeratin (34BE12), CD3, CD14, CD68, and CD163. Whole slides were imaged with the Zeiss Axioscanner 7.0 and analyzed with VIS from Visiopharm (v2023.11). The tumor regions were initially detected by an AI Deeplab algorithm based on the R6G signal. In tumor and non-tumor regions, nuclei and cytoplasm were detected by a second AI U-net algorithm based on the DAPI signal. Next, individual fluorescent channels were quantified based on the median intensities. Downstream processing was done in R with SPIAT (v1.0.4.). Cells were annotated according to positivity of the following channels: Cytokeratin+CD- (Tumor); CD14+CD3-CD68-CD163- (Monocyte); CD68+ (Macrophage); CD163+CD68+ (M2 Macrophage); CD3+ (T-cell). Following annotation, cells were visualized based on the center coordinates of their nuclei. Cell type fractions were computed for each core and cohort-wide statistics were performed with the ggpubr package (v0.4.0).

### Bulk RNAseq and bioinformatic data analysis

Cell lines were grown to 60-70% confluency before RNA extraction was performed with TRIzol™. A subsequent DNAse treatment was done with the TURBO DNA-free Kit (Invitrogen) to purify the samples. Next, samples were sequenced with the DNA nanoball (DNB) sequencing protocol (BGI) to a depth of 50M reads/sample. The SOAPnuke pipeline (BGI) was used to perform quality checks and preprocessing. Clean reads were mapped to the human reference genome (hg19) with the STAR aligner (v2.7.9a) and the GENCODE v35 annotation. Duplicates were marked with Sambamba (0.8.0) and gene counting was done with FeatureCounts (v2.0.0). Downstream analysis was performed in R using the DESeq2 package (v1.36.0). After variance stabilizing transformation, a principal component analysis was performed using the top500 variable expressed genes. Differentially expressed genes were visualized with the Enhanced volcano package (v.1.14.0). Gene ontology and enrichment analysis were computed based on the Hallmark gene sets from the molecular signature database using the enrichR (v.3.1) and the fgsea (v.1.22.0) packages, respectively.

### Chip Sequencing data analysis

CHD4 chromatin binding peaks from ChIP-sequencing of the SCC9 cell line, overexpressing CDK2AP1 or transfected with an empty vector, were retrieved from (18). Bedops (65) was utilized to identify consensus peaks for each condition by selecting peaks identified in at least two replicates. Peaks were then annotated using the ChIPseeker (version 1.30.3) R-package (66). Over-representation analysis was performed on CHD4 binding peaks present exclusively in the CDK2AP1 overexpressing samples through the clusterProfiler (version 4.2.2) R package(7) using the MSigDB Hallmarks (67) annotation.

### Data availability

RNA sequencing data regarding the gene expression profiling of oral squamous cell carcinoma (OSCC) cell lines either Parental or CDK2AP1 knockouts (KO) has been deposited in the gene expression omnibus and is publicly accessible with identifier GSE260789. Gene expression profiling of the CA1 cell line w/wo silencing of BRM and BRG1 can be accessed with identifier GSE260831.

### Statistical analysis

Results are shown as means of three biological replicates, with error bars representing the standard error of the mean (S.E.M). Details of each statistical test are indicated in the figure legends. All statistical tests and graphs were executed using the R software package.

### Ethics Statement

Human tissues and patient data were used according to “The Code of Conduct for Responsible Use” and “The Code of Conduct for Health Research” as stated by the Federation of Dutch Medical Scientific Societies (20). Furthermore, The Erasmus MC Medical Ethics Committee approved the research protocol (MEC-2016-751).

## Acknowledgments

We thank all members from the Fodde laboratory for helpful discussions. We are grateful to the ErasmusMC for proving support to this study through the Mrace grant 2027.Furthermore, Adrian Biddle is a member of the Barts Centre for Squamous Cancer, funded by Barts Charity.

## Author contributions

RS and RF conceived the experiments and wrote the manuscript. R.S, C. E, M.P.V, F.A.T., R.J. performed experiments. S.K., B. vd S., M.J. de H, J.A and R. B de J. advised on the experimental strategy and provided the clinical material. T.vd B. and A.L.N. performed the TMA multiplex staining and initial computational analysis. P.L provided useful insights in the field of immunology and together with R.S. and R.F. wrote the manuscript. R.F., in collaboration with P.V. supervised the study and was responsible for concept and design of the study.

---

**Supplementary Figure 1.**
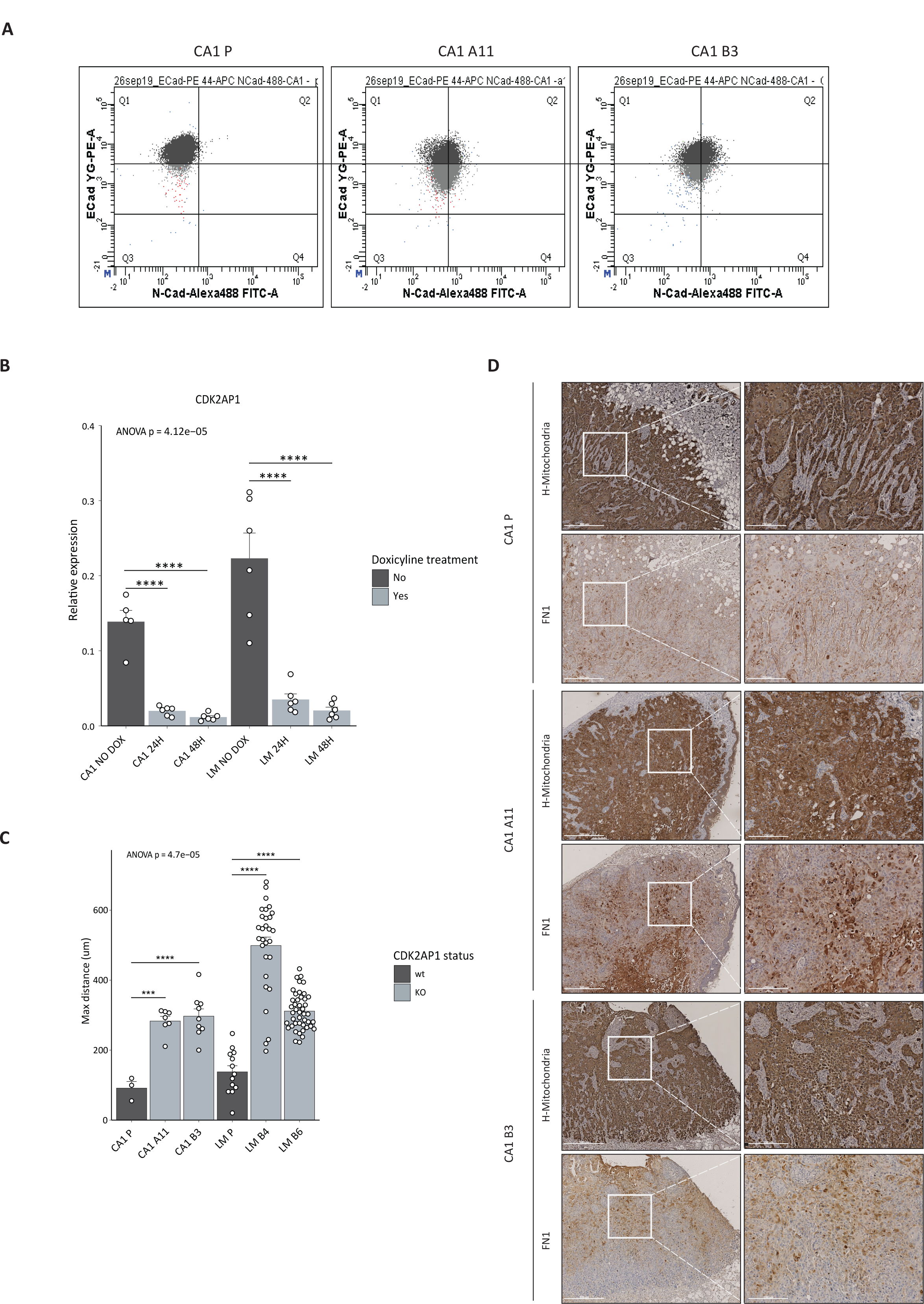
CDK2AP1-KO confers epithelial mesenchymal plasticity in OSCC cell lines. **A.** Flow cytometric analysis of the CA1 parental and *CDK2AP1*-KO cell lines (CA1-A11 and -B3) with antibodies directed against E-CAD and N-CAD. E-CAD/N-CAD-low and -high regions were defined using the parental cell line as a reference. **B.** *CDK2AP1* RT-qPCR expression analysis upon shRNA induction in parental CA1 and LM cell lines. Experiments were performed in triplicate and mRNA expression was normalized to that of GAPDH. p-values denote one-way ANOVA and one-sample t-test against the uninduced parental cell lines (*p<0.05; **p<0.01; ***p<0.001). **C.** Analysis of the maximal distance covered by the parental and *CDK2AP1*-KO CA1 and LM cell lines in the invasion assay (see lower panel in Fig. 1C). Each dot represents a single-cell measurement. Experiments were performed in triplicate and p-values denote one-way ANOVA and one-sample t-test against the parental cell lines (*p<0.05; **p<0.01; ***p<0.001). **D.** IHC analysis of tumors obtained by subcutaneous injection of either CA1 parental or *CDK2AP1*-KO clones (CA1-A11 and -B3) with antibodies against human mitochondria and FN1. In the left column, lower magnification (4X) images are shown a specific area of which is shown at higher magnification (20X) in the right column. Lower magnification scale bars: 500 μm, higher magnification field scale bars: 250 μm.

**Supplementary Figure 2.**
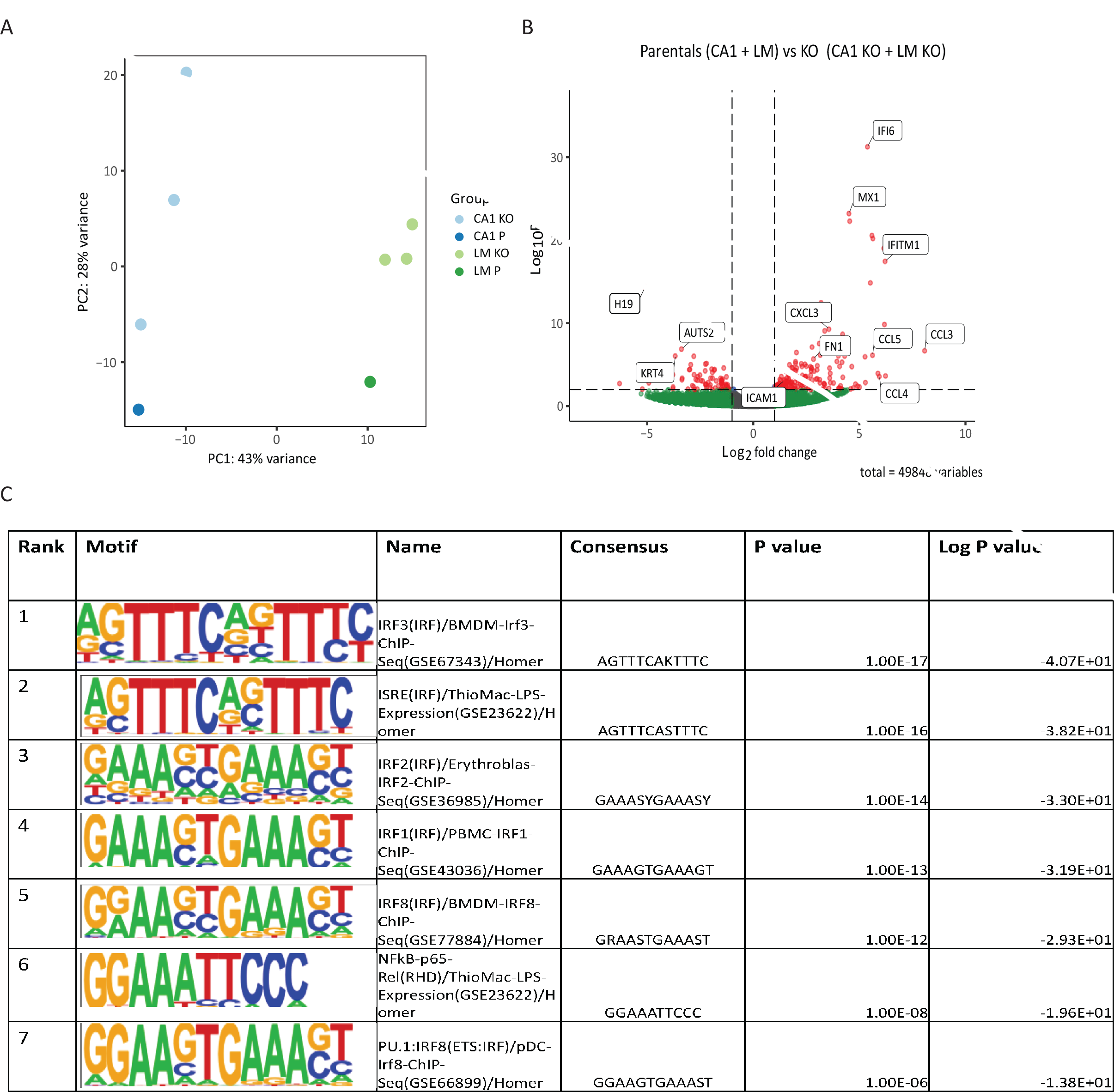
RNAseq analysis of parental and CDK2AP1-KO OSCC cancer cell lines reveals the downstreamregulation of the inflammatory pathways. **A.** Principal component analysis (PCA) of RNAseq profiles from parental and *CDK2AP1*-KO OSCC cancer cell lines. **B.** Volcano plots of the differentially expressed genes upon *CDK2AP1* ablation in our panel of OSCC cell lines (log_2_ fold change> 2). On the right side of the plot, genes upregulated upon deletion of CDK2AP1 in CA1 and LM -KO clones are shown; while on the left side, the genes downregulated after CDK2AP1 KO are represented. **C.** HOMER transcription factor motive analysis of the differentially expressed genes upon CDK2AP1 ablation in the CA1 and LM OSCC cell lines.

**Supplementary Figure 3.**
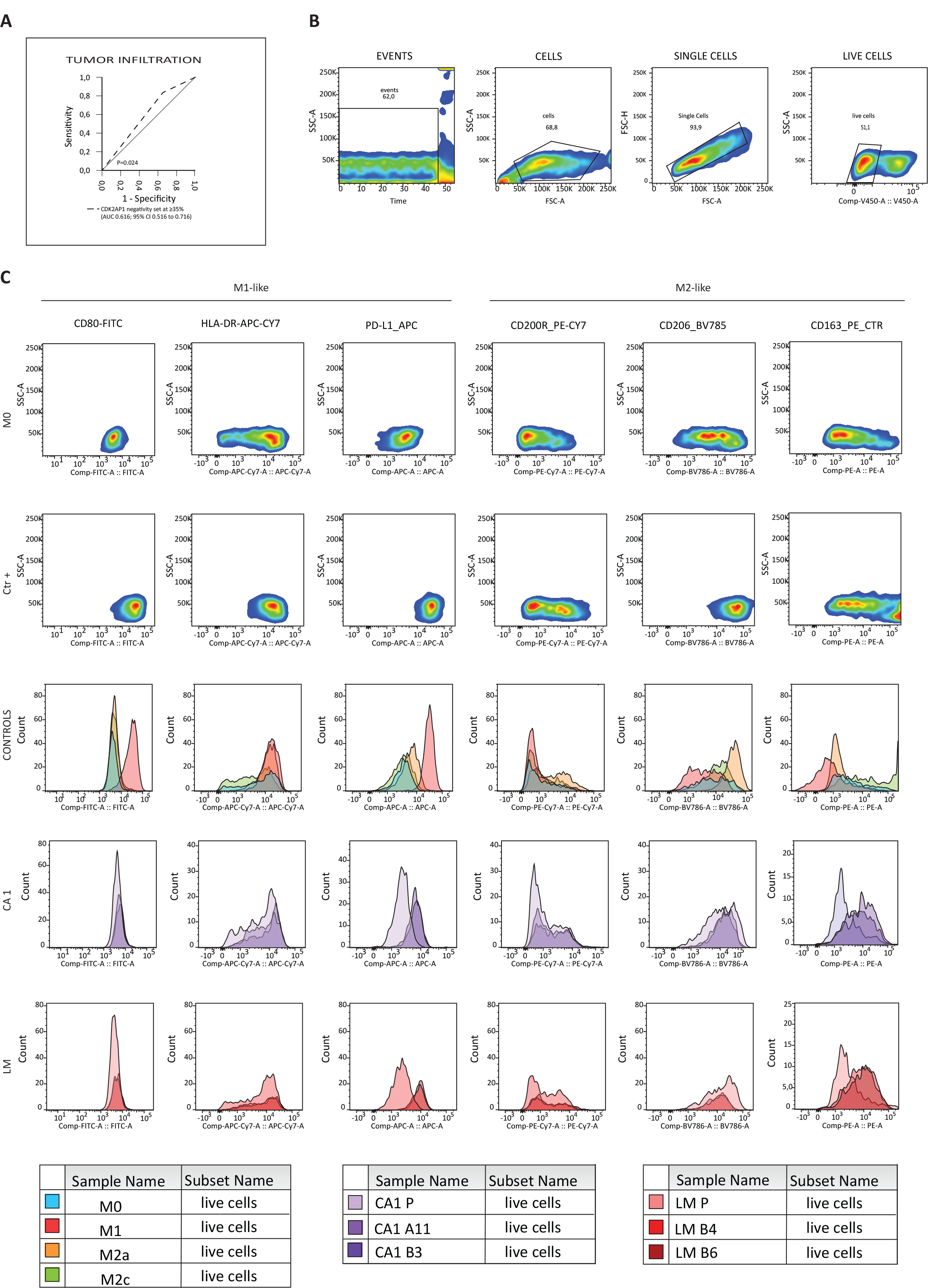
CDK2AP1 status influences the immune cells of the tumor microenvironment (TME) **B.** The ROC (receiver operating characteristic) curve based on which the optimal threshold of 35.0% of CDK2AP1-negative cells was established and employed for tumor infiltration analysis. **C.** Cytofluorimetric pre-processing strategy for the detection of the different states of macrophage polarization. Briefly, the number of cells were determined based on cell size over time (EVENTS). Afterwards, using FSC-A and FCS-H the cells were firstly separated by the debris (CELLS) and then the single cells were purified (SINGLE CELLS). Eventually, thanks to the ZombieViolet staining, we could focus on the living cells (LIVE CELLS) (see also Materials and Methods) Expression of markers of macrophages polarization by cytometric bead assay. The first lane shows the profile of each marker detected in our negative control sample (M0 macrophages) while the second lane shows the profile of the positive controls (Ctr +): for the M1-like specific markers, the M1-like macrophages treated with INF-γ + LPS are shown; for M2-like macrophages, the ones treated with IL-4 and IL-10 are reported. The third lane shows the different florescence levels of the controls taken in exams, per marker (CONTROLS). The last two lanes show the different profiles obtained in the case where the macrophages were stimulated with the supernatant of the CA1 and LM cells lines respectively (Parental profiles in light colors; CDK2AP1 KO clones in dark colors).

